# *In vivo* production of an anti-HIV antibody from primate hematopoietic cells by non-viral knock-in

**DOI:** 10.1101/2025.05.02.651933

**Authors:** Jack M.P. Castelli, Katrina Poljakov, Youngseo Jwa, Molly E. Cassidy, Matthew D. Gray, Jocelyn N. Sanchez Gaytan, Mark R. Enstrom, Jonathan D. Linton, Anthony Rongvaux, Justin J. Taylor, Jennifer E. Adair

## Abstract

Non-viral gene editing offers a practical alternative to viral delivery for durable biologics production. Clinical trials have shown that adeno-associated virus encoding broadly neutralizing antibodies can protect against HIV, but result in limited, short-lived responses. The development of non-viral gene editing approaches in hematopoietic stem and progenitor cells holds promise for long-term antibody production. In this study, we evaluated CRISPR/Cas9 and CRISPR/Cas12a for gene knock-in at the immunoglobulin heavy chain locus in non-human primate hematopoietic stem and progenitor cells. Delivering the nuclease as a protein alongside a custom DNA template, we optimized editing with Cas12a and demonstrated higher knock-in efficiency and fewer non-specific edits than Cas9. Transplantation of edited non-human primate hematopoietic stem and progenitor cells into MISTRG mice led to engraftment, B cell differentiation, and transgene expression of a reporter transgene or anti-HIV antibody after HIV immunization with detectable anti-HIV antibody titers in peripheral blood circulation. These findings demonstrate the feasibility of using non-viral gene editing in HSPC as a potential strategy for sustained biologics production in the treatment of chronic diseases such as HIV. Future work will assess the efficacy of this model in a non-human primate model of HIV infection.

## INTRODUCTION

Broadly neutralizing antibodies (bnAb) against human immunodeficiency virus (HIV) isolated from long-term non-progressors are capable of providing immune protection against diverse strains of HIV-1.^1^ They have been demonstrated as viable treatment alternatives to controlling viral loads in non-human primate (NHP) models and in human clinical trials.^2,3^ As opposed to standard-of-care antiretroviral therapy (ART), bnAbs can be administered less frequently and sustain activity in circulation for up to 6 months, a key advantage for reaching vulnerable populations who cannot achieve daily adherence or suffer from ART-associated long-term toxicities.^4^ Previous studies have explored the delivery of bnAb transgenes to various cell types using viral vectors.^5,6^ When delivered into muscle tissues in animal models, adeno-associated viral (AAV) vectors can achieve bnAb expression and protect from viral challenges.^7,8^ However, this indiscriminate production of antibody, combined with the immunogenicity of AAV, results in host anti-antibody and anti-AAV responses.^9–11^ These responses curtail the sustained production of bnAbs and prevent re-treatment. Immunization strategies have also failed to induce endogenous bnAb expression, in part due to extensive somatic hypermutation required to attain high-affinity bnAb sequences, as well as poly- and auto-reactive precursor sequences that fall prey to negative selection during maturation of germinal center B cells.^12–15^

CRISPR/Cas technology has enabled precise targeting of the immunoglobulin heavy chain (*IGH*) locus to replace endogenous antibodies with transgenes delivered by viral vectors.^16–23^ Edited B cells expressing engineered antibodies have conferred anti-viral protection in murine models.^16,17,20^ It remains unclear how long such gene-edited B cells can sustain an immune response, though those that develop into long-lived plasma cells could confer immune memory.^19,20^ Notably, one of the earliest cell types explored for bnAb transduction were hematopoietic stem and progenitor cells (HSPC), the precursors to all B cells.^24^ These lentivirally-transduced HSPCs could subsequentially differentiate into bnAb-producing plasma cells *in vitro* but antibody production was driven from proviral elements semi-randomly integrated into the genome.^24^ If the endogenous antibody-producing *IGH* locus of HSPC could be precisely edited non-virally and retain the ability to engraft and differentiate into antibody-producing B cells that retain the antibody transgene *in vivo*, this approach could provide a durable source of bnAb-producing B cells without viral vector-triggered immune responses. While *in vivo* delivery of CRISPR/Cas to HSPC with high specificity remains a challenge, animal models offer valuable insights into hematopoiesis and engraftment potential.

Here, we utilized the MISTRG mouse strain (*CSF1*^h/h^ *IL-3*/*CSF2*^h/h^*SIRPA*^tg^ *THPO*^h/h^ *Rag2*^−/−^ *Il2rg*^−/−^), which supports the engraftment of NHP HSPC, to investigate the development of edited HSPC and transgene expression following non-viral knock-in.^25,26^ The use of NHP cells enables the development of editing materials, dependent on genomic context, for a relevant pre-clinical model of HIV infection. Our findings in this primatized mouse model demonstrate the potential for durable bnAb production from hematopoietic cells as a strategy for controlling HIV.

## RESULTS

### Optimized editing of *IGH* locus in primate HSPC

We first characterized and tested CRISPR guide RNA (gRNA) for targeted gene knock-in in NHP HSPC (CD34^+^). Sequence data for the *IGH* locus was available for rhesus macaques (*Macaca mulatta*) but had not been documented for pigtail macaques (*M. nemestrina*), a closely related and larger species that has been used as a model for HIV-1 infection and long-term HSPC engraftment studies.^27,28^ We sequence-verified the region of this locus spanning from the final J segment to the Eμ enhancer in *M. nemestrina*, an intronic region with low heterogeneity that is unchanged by variable (V), diversity (D) and joining (J) segment (VDJ) recombination (Figure 1A).^29^ Using the verified sequence, we screened for all potential CRISPR/Cas9 and CRISPR/Cas12a guide RNA (gRNA) by identifying relevant protospacer-adjacent motif (PAM) sequences within this locus. Cas9 and Cas12a nucleases generate site-specific DNA cuts but with different free DNA end configurations and with different PAM sequences, with the latter associated with higher specificity.^30^ Chen *et al*. demonstrated that sequence context can influence the outcome of gene editing following CRISPR treatment, with Tatiossian *et al*. showing that larger deletions are associated with an increased likelihood of homology-directed repair (HDR) at a specific locus.^31,32^ For these reasons, a total of 120 Cas9 gRNA and 56 Cas12a gRNA were identified and ranked based on predicted efficiency and accuracy using the CRISPOR webtool (https://crispor.gi.ucsc.edu/, Table S1).^33^ Lindel (logistic regression model to predict insertions and deletions) scores were included as a measure of HDR pathway preference following double-strand breaks induced by Cas9 to minimize the production of undesired mutations with these gRNA.^31^ We then synthesized the top five gRNA for each CRISPR/Cas system for empirical testing.

**Figure 1.**
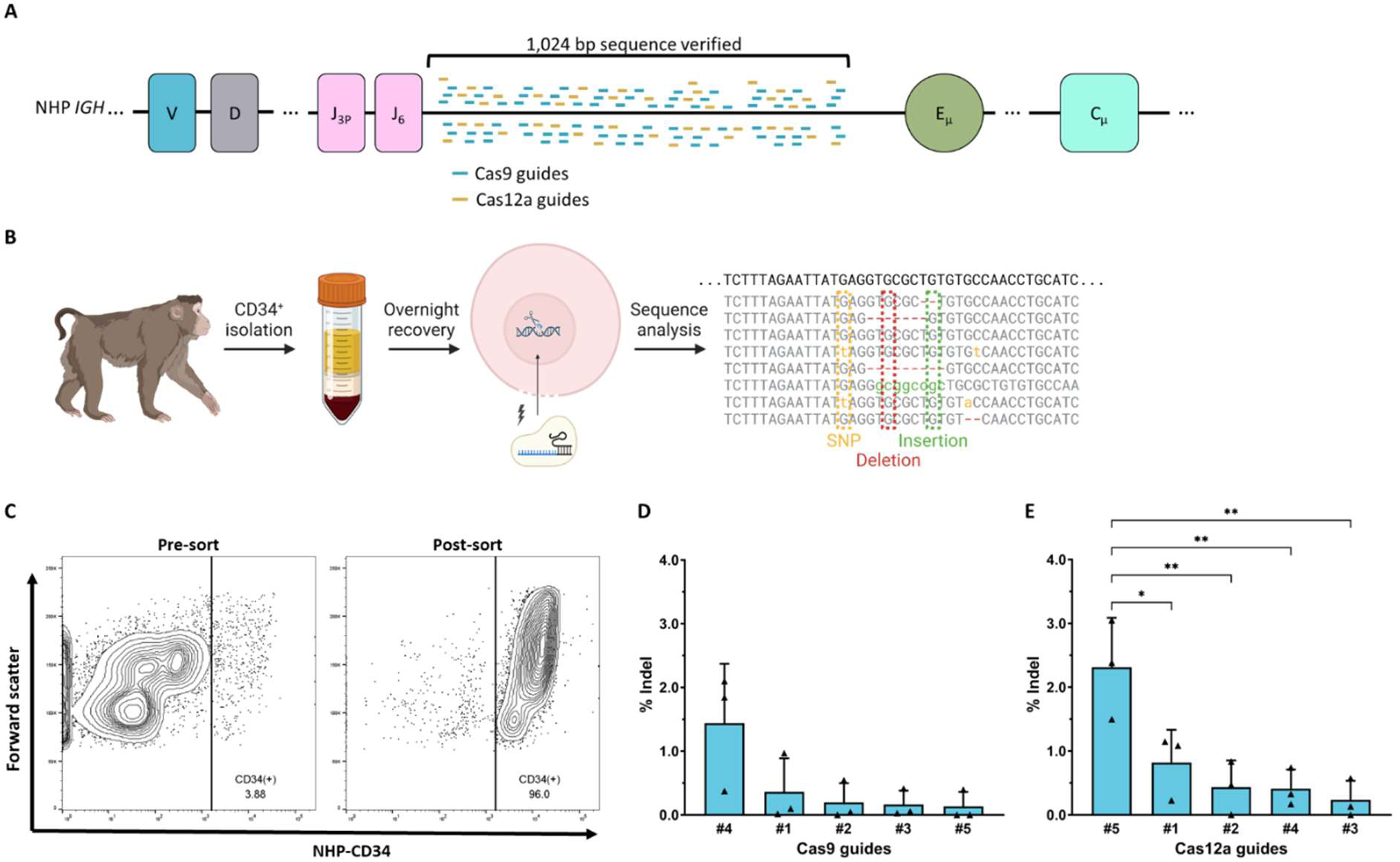
CRISPR/Cas guides capable of editing of *IGH* locus in NHP HSPC. (A) Target genomic site within the NHP *IGH* locus, between the *VDJ* genes and the constant regions, sequence verified in two NHP species, *M. mulatta* and *M. nemestrina*. All possible Cas9 and Cas12a guides were screened within this sequence. (B) CD34^+^ HSPC were isolated from *M. nemestrina* bone marrow aspirates and electroporated with RNP. Target locus was analyzed for genetic edits, including single nucleotide polymorphisms (SNP), insertions, and deletions (indel). (C) CD34^+^ purity following immunomagnetic separation as measured by flow cytometry. (D, E) Indel frequency of selected Cas9 (D) and Cas12a (E) guides 3 days after electroporation of 10^6^ CD34^+^ cells (n = 3 biological replicates, 3 donors). Data are presented as mean ± standard deviation (SD). Significance was calculated with an ordinary one-way ANOVA with *p<0.05, and **p<0.01.

To evaluate the performance of these guides, we isolated primary bone marrow-derived CD34^+^ hematopoietic stem and progenitor cells (HSPC) from healthy adult NHP donors (Figure 1B). CD34^+^ cells were purified from bone marrow aspirates, reaching 79-97% purity post-enrichment (Figure 1C, Table S2). Cells were electroporated with CRISPR ribonucleoproteins (RNP), and insertion and deletion (indel) formation at the target loci was assessed through high-throughput sequencing. Cas9 gRNA exhibited indels in up to 1.4% of sequence reads (Figure 1D). Cas12a gRNA demonstrated higher average editing levels, albeit without statistical significance (0.84% ± 0.38% vs. 0.46% ± 0.25%, unpaired two-tailed t-test, p = 0.42), with the most efficient gRNA achieving a 2.3% indel frequency (Figure 1E). This was statistically higher than other gRNA (ordinary one-way ANOVA, p = 0.0026), although overall editing efficiencies were low. These results demonstrate feasibility of gene editing in NHP HSPC, with Cas12a showing a slight advantage in editing efficiency over Cas9 at this locus.

### Achieving HDR at *IGH* locus and transgene expression *in vitro*

In addition to sequence context which can influence HDR frequency, we assessed the ability of CRISPR/Cas9 and CRISPR/Cas12a systems to mediate the knock-in of a DNA template into the target *IGH* locus via HDR. Cas9 nuclease generates a blunt double-strand break, while Cas12a introduces staggered cuts with 5 base pair (bp) single-stranded overhangs.^34^ These differences in DNA cleavage can also influence HDR outcomes, resulting in either productive insertion of the DNA template at the correct locus with proper transgene expression or non-productive insertions that are truncated, duplicated, or introduce additional indels that may compromise transgene expression (Figure 2A).

**Figure 2.**
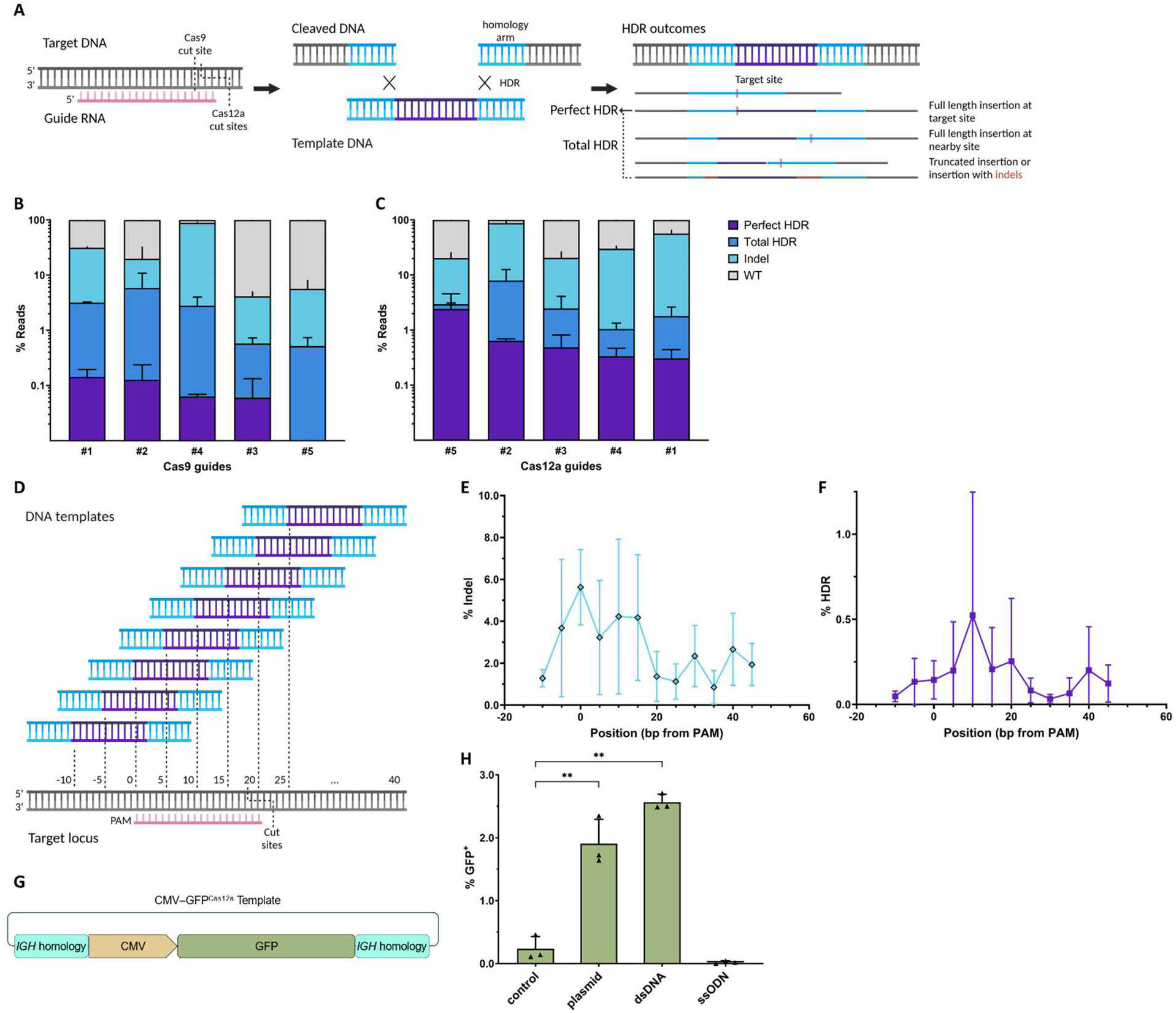
Guides associated with HDR events using short templates enable knock-in and expression of transgenes encoding cassettes. (A) Cas9 and Cas12a guides produce distinct cuts, allowing knock-in of template with homology to cut site, and result in different HDR outcomes. (B, C) HDR frequency with Cas9 (B) and Cas12a (C) guides following electroporation in 10^6^ LCL 8664 cells, a cell line derived from *M. mulatta* lymphocytes (n=3 experimental replicates). Reads were classified by editing outcomes. (D) ssODN templates positioned for insertion at different distances from PAM, varying from −10 bp to +40 bp. (E, F) Fraction of reads containing indel and perfect HDR events following electroporation in 10^6^ LCL 8664 cells (n = 3 experimental replicates). (G) GFP template was constructed for knock-in and genomic expression. Homology arms were designed for insertion at cut site of optimal Cas12a guide. (H) Stable GFP expression 15 days following electroporation in 10^6^ LCL 8664 cells using different templates: plasmid without RNP (control), plasmid, dsDNA, or ssODN with RNP (n = 3 experimental replicates). Data are presented as mean ± SD, significance calculated with an ordinary one-way ANOVA.

To quantify HDR efficiency and fidelity, we electroporated a NHP B cell line (LCL 8664) with ribonucleoproteins (RNP) and a small single-stranded oligodeoxynucleotide (ssODN) template. Sequencing of the edited cells allowed us to classify genotypes based on productive or non-productive HDR events. Cas9 gRNA achieved up to 5.9% total HDR, with nearly half of these events being non-productive (Figure 2B). In contrast, Cas12a gRNA exhibited a higher total HDR efficiency, reaching up to 8.0%, and showed a much lower fraction of non-productive HDR events (Figure 2C). Based on these findings, we selected a Cas12a gRNA that showed a favorable ratio of productive HDR to non-productive HDR and indels for further experiments (NHP *IGH* Cas12a guide #5: TTTAGAATTATGAGGTGCGC). To further optimize HDR efficiency, we designed and tested ssODN templates positioned at different distances from the predicted cut site of this optimal Cas12a gRNA (Figure 2D).^35^ Among the tested configurations, HDR templates placed +10-20 bp downstream from the first gRNA base exhibited the highest HDR efficiency, along with the highest ratio of productive HDR to indels (Figure 2E). Subsequent templates were designed with insert sequences placed +15 bp from the Cas12a gRNA.

We next designed a large 2,250 bp DNA template encoding a green fluorescent protein (GFP) reporter transgene under the control of a constitutive cytomegalovirus (CMV) promoter, flanked by 450 bp homology arms aligned to the NHP *IGH* locus (Figure 2F). Three versions of this CMV-GFP^Cas12a^ template were tested for their ability to induce GFP expression when delivered by electroporation to LCL 8664 cells alongside Cas12a RNP including our optimal gRNA. GFP expression was monitored over several cell divisions. LCL 8664 cells electroporated with double-stranded DNA (dsDNA) or plasmid templates with Cas12a RNP showed durable GFP expression, indicating successful knock-in of the transgene (Figure 2G). The dsDNA template produced higher GFP expression than plasmid template alone or with Cas12a RNP, indicating greater efficiency for transgene knock-in. In contrast, co-delivery of ssODN template was not associated with significant GFP expression. Thus, we proceeded with dsDNA templates for our non-viral editing approach in mouse studies.

### Engraftment and hematopoiesis of edited NHP HSPC in MISTRG mice

We next evaluated the potential for transgene knock-in, engraftment, and differentiation potential of primary CD34^+^ NHP HSPC. To achieve this, we utilized the immunodeficient MISTRG mouse model, which supports NHP cell engraftment.^26^ NHP HSPC were electroporated with Cas12a RNP and a 2,384 bp VH4-GFP^Cas12a^ construct expressing GFP under the control of a B cell-specific promoter derived from NHP variable heavy (VH) domain allele 4-38-2, the most conserved allele and most abundant VH transcript.^36^ Electroporated HSPC were then transplanted into sub-lethally irradiated, neonatal MISTRG mice (Figure 3A). Engraftment and development of NHP cells were tracked through regular peripheral blood draws and in different tissues at necropsy.

**Figure 3.**
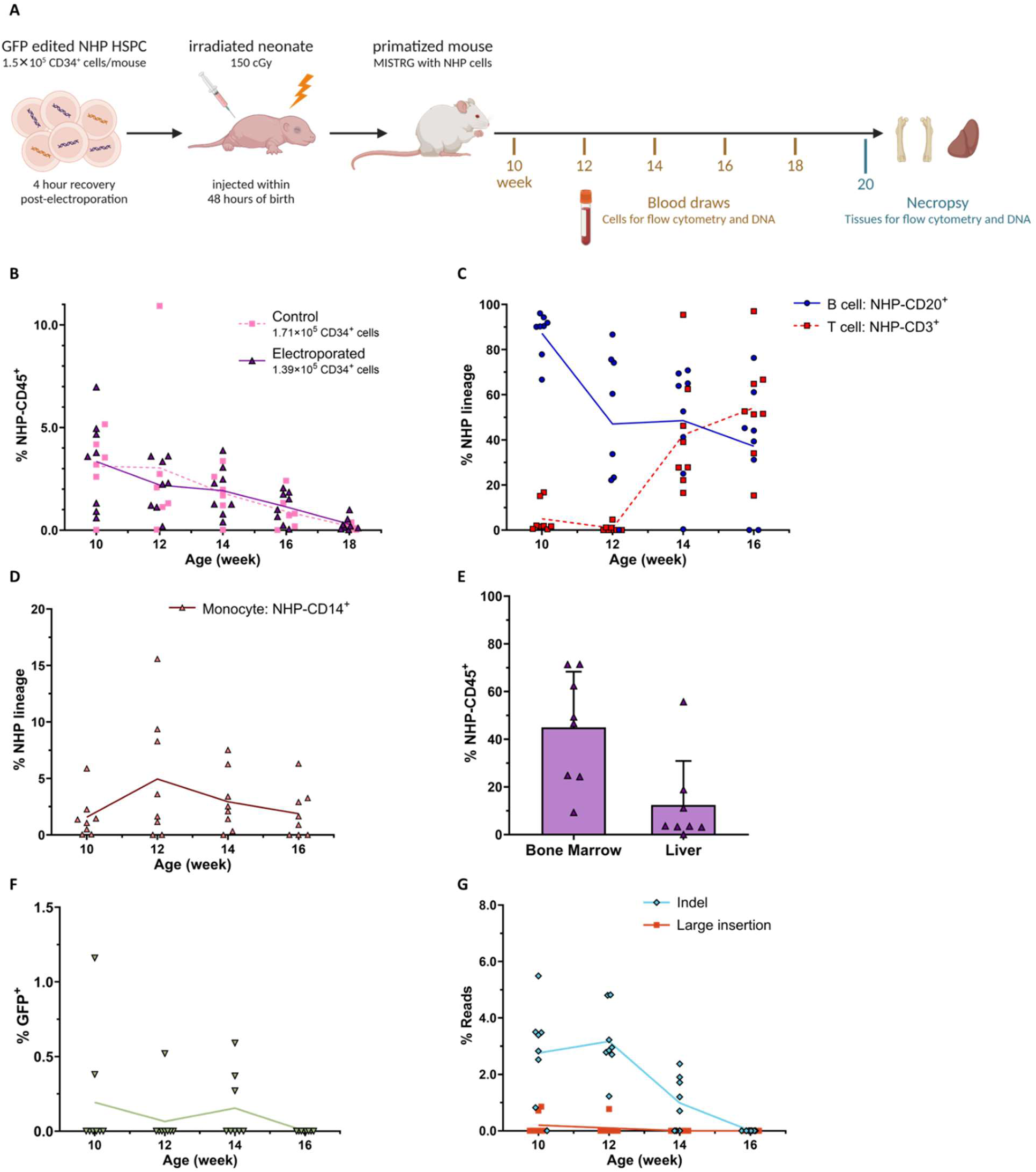
Edited HSPC engraft in MISTRG model and express inserted transgene. (A) NHP HSPC electroporated with VH4-GFP^Cas12a^ were injected into sub-lethally irradiated, neonatal MISTRG mice. Hematopoietic lineages in the primatized mice were monitored over time and at necropsy. (B) Engraftment levels in peripheral blood over time as measured by flow cytometry on NHP-CD45^+^ cells from mice injected with electroporated or control HSPC. n = 8 mice, n = 2 NHP donors (electroporated group, 1.39 × 10^5^ CD34^+^ cells/mouse). n = 6 mice, n = 2 NHP donors (control group, 1.71 × 10^5^ CD34^+^ cells/mouse). (C, D) Multilineage engraftment in peripheral blood over time as measured by frequency of B cells (CD3^-^ CD20^+^), T cells (CD3^+^ CD20^-^), and monocytes (CD3^-^ CD20^-^ CD14^+^) in NHP-derived populations (NHP-CD45^+^). (E) Engraftment levels in the bone marrow (femur) and the liver at necropsy, performed at week 20. Error bars represent SD. (F) *In vivo* transgene expression as measured by frequency of GFP^+^ cells in NHP-derived populations (NHP-CD45^+^), as compared to total B cell frequency in peripheral blood over time. (G) Editing levels at the NHP *IGH* locus in peripheral blood as measured by high-throughput sequencing on MiSeq platform. Large insertions represented by insertions >8 bp. Lines represent means in (B, C, D, F, and G). n = 8 mice, n = 2 NHP donors (electroporated group, week 18 omitted for low engraftment levels, C-G).

Results showed that electroporated NHP HSPC were able to successfully engraft in immunodeficient MISTRG mice. Mean NHP-CD45^+^ white blood cell (WBC) populations in peripheral blood reached up to 3.3% at peak engraftment (Figure 3B). Increased HSPC cell doses trended towards higher engraftment (Figure S1). Engraftment persisted for over 16 weeks without visual signs of graft-versus-host disease (GVHD). To assess multilineage differentiation, mice injected with 1.4 × 10^5^ VH4-GFP^Cas12a^ electroporated HSPC were monitored for white blood cell populations, including lymphocytes and monocytes. Early after transplantation, we observed mostly B cell populations, with T cell populations emerging around week 12, eventually surpassing B cells by week 16 (Figure 3C). Monocyte levels remained consistent and low throughout the study (Figure 3D). Hematopoietic NHP cells were detected in the bone marrow and liver at necropsy (Figure 3E). These trends were also observed in MISTRG mice receiving control (i.e., non-edited) HSPC (Figure S3).

GFP expression was detected in 50% of mice transplanted with edited HSPC, with four out of eight mice showing GFP^+^ cells in peripheral blood. From weeks 10 to 14, 0.41% to 0.77% of NHP-derived cells expressed GFP (Figure 3F). However, a decline in GFP expression coincided with decreasing B cell populations and engraftment levels. Further sequencing confirmed the presence of edited cells *in vivo* (Figure 3G). Indels, including large insertions, were detected at the target NHP *IGH* locus in the early weeks post-transplantation but had drastically declined by week 16 when peripheral blood B cells also declined.

### *In vivo* expression of a bnAb from the *IGH* locus in primate B cells

We next developed templates for expressing a bnAb transgene. Our 2,790 bp template design was based on previous efforts to engineer antibody knock-in at the human *IGH* locus.^29^ Specifically, we constructed a sequence containing the VH4 B cell-specific promoter, along with a complete light chain and recombined heavy chain separated by a short linker, followed by a splice site allowing the integration of an endogenous heavy chain constant region (Figure 4A). The coding sequences for bnAb 10-1074, which targets the V3 glycan site on the HIV envelope protein, were optimized for NHP expression, resulting in the VH4-10-1074^Cas12a^ template.

**Figure 4.**
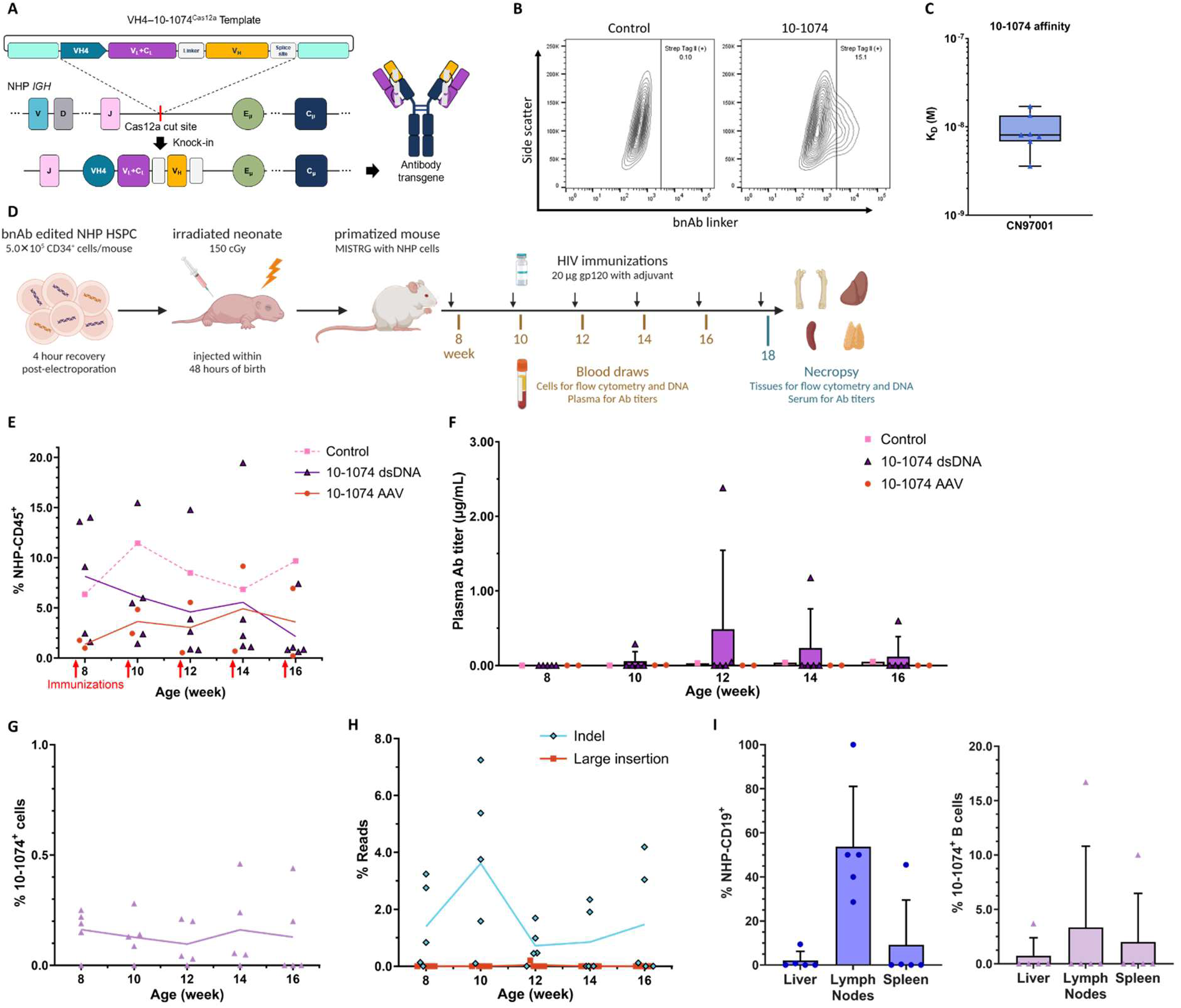
bnAb transgene expression and immunization in MISTRG mice. (A) bnAb template was designed for expressing the full light chain and the variable heavy chain fragment of 10-1074. Following productive knock-in at the target locus, bnAb expression replaces endogenous antibody production, bypassing VDJ recombination and forming a full-length transcript via splicing with a constant heavy chain. (B) *In vitro* 10-1074 production as confirmed by staining of cell surface transgene linker. Flow cytometry was performed on HEK 293E cells following transduction with an expression vector containing an IgG constant heavy chain. (C) Binding of recombinant bnAb to HIV antigen, the gp120 protein of strain CN97001, as measured by bio-layer interferometry (BLI) and quantified as the dissociation constant (K_D_). n = 6 technical replicates. (D) NHP HSPC were electroporated with VH4-10-1074^Cas12a^, either as dsDNA or AAV, and transplanted into MISTRG mice as before. Beginning at week 8, mice were immunized with CN97001 gp120 injections once every two weeks until necropsy. Control mice were injected with unedited HSPC. (E) Engraftment levels in peripheral blood over time as measured by flow cytometry on NHP-CD45^+^ cells in mice injected with edited HSPC using different templates. (F) *In vivo* anti-HIV antibody titers were measured by ELISA with a biotinylated gp120 on diluted plasma samples. dsDNA: n = 5 mice, n = 2 NHP donors. AAV: n = 2 mice, n = 2 NHP donors. Control: n = 1 mouse, n = 2 NHP donors. n = 2 technical replicates per sample. (G) Cell surface expression of 10-1074 in peripheral blood as measured by flow cytometry with antibody against its linker. (H) Editing levels at the NHP *IGH* locus in peripheral blood as measured by high-throughput sequencing (MiSeq). Large insertions represent insertions >8 bp. (I) B cell populations and fraction of B cell population expressing 10-1074 at necropsy, week 18, as measured by flow cytometry on NHP-CD19^+^. n = 5 mice, n = 2 NHP donors (10-1074 dsDNA group, G-J). Error bars represent SD.

To validate expression of VH4-10-1074^Cas12a^, we transfected HEK 293E cells *in vitro* with plasmid and measured transgene surface expression levels (Figure 4B). Recombinant bnAb was isolated from cell culture supernatant and subsequently assessed for binding affinity to an HIV envelope protein antigen. Using gp120 monomers from various HIV strains, we observed that HIV strain CN97001 exhibited the highest affinity for recombinant 10-1074 (Figure 4C, Figure S3). These results confirmed that the transgene design permitted 10-1074 binding to its target antigen.

In addition to a dsDNA template amplified from the plasmid, we produced an AAV template to compare non-viral and viral knock-in strategies for NHP HSPC. We compared a panel of AAV serotypes to transduce NHP HSPC and selected AAV6.2 serotype for experiments (Figure S4). Subsequently, NHP HSPC edited with each of these VH4-10-1074^Cas12a^ templates were injected at a dose of 5.0 × 10^5^ cells into MISTRG mice (Figure 4D). To promote clonal expansion of B cells with productive bnAb knock-in, beginning at 8 weeks of age, mice were immunized with HIV-CN97001 gp120 monomers. Peripheral blood and plasma samples were collected to monitor lymphocyte development and antibody production. Multiple tissues, including secondary lymphoid organs (lymph nodes, spleen), bone marrow, and liver, were collected at necropsy for further analysis.

As with previous experiments, edited NHP HSPC demonstrated successful engraftment and differentiation into mature cell types, including large B cell populations early in the study (Figure 4E, Figure S5). With CD34^+^ doses of 5.0 × 10^5^, up to 8.2% NHP-CD45^+^ cells were observed at peak engraftment in peripheral blood of mice receiving dsDNA templates. Four weeks after immunization, anti-HIV antibody titers were detectable in a subset of primatized mice (Figure 4F).

In the cohort receiving the dsDNA template, one of five (20%) mice developed antibody titers up to 2.4 µg/mL. 10-1074-expressing cells were identified in all primatized mice, although cell-surface expression was low, with less than 0.46% of NHP-derived cells observed in peripheral blood (Figure 4G). Interestingly, the mouse with detectable antibody titers in plasma had the highest NHP CD45^+^ WBC engraftment levels but did not show a significantly larger population of 10-1074-producing cells, suggesting that most antibody-expressing cells were likely not in circulation (Figure S6). Hematopoietic NHP cells were detected in all tissues analyzed at necropsy, including CD34^+^ cells in the bone marrow (Figure S7).

Mean indel frequency peaked at 1.5% in peripheral blood at week 16 (Figure 4H). Sequencing of cells from bone marrow, liver, spleen, and lymph nodes confirmed the continued presence of edited cells in different tissues at necropsy (Figure S6). While antibody production was not significantly correlated with indel levels, large insertions (defined as ≥8bp, 0.20% of sequence reads) were observed in the mouse with detectable antibody titers. Lymphocyte populations followed the same trends as previously observed (Figure S5). B cell populations declined with similar kinetics as previous experiments despite immunization. The mouse with detectable antibody titers demonstrated a decline in B cell populations, with T cell populations taking over at week 12 (Figure S7). NHP-CD19^+^ B cells were present in the lymph nodes of all mice, as well as the liver and the spleen of a subset of mice at necropsy, where B cell maturation occurs in germinal centers, of which up to 17% were 10-1074-producing cells (Figure 4I).

### Antibody expression and B cell maturation in MISTRG6 mice

To further validate *in vivo* 10-1074 production, we repeated the injection of edited NHP HSPC in MISTRG6 mice, a strain derived from MISTRG with the additional knock-in of human interleukin-6 (IL-6).^37^ Given IL-6’s role in promoting B cell development and immunoglobulin G (IgG) production, we hypothesized that antibody titers would be detectable in a larger fraction of animals. To compare viral and non-viral editing conditions, seven mice were injected with HSPC receiving VH4-10-1074^Cas12a^ as either dsDNA or AAV6.2 template (Figure 5A). In addition, tissue analysis was performed at an earlier timepoint closer to peak antibody expression to identify the location of antibody-producing cells.

**Figure 5.**
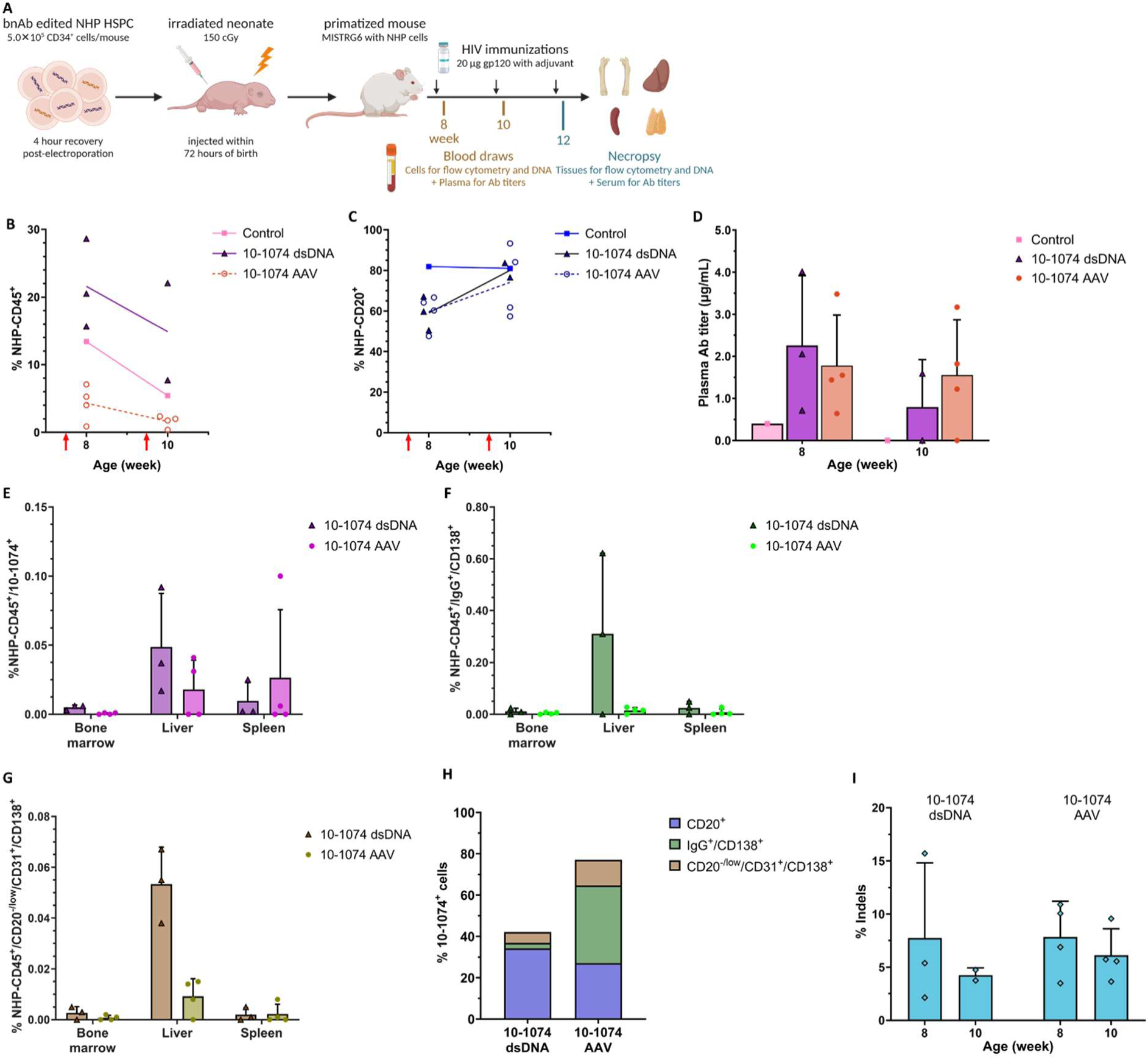
MISTRG6 mice support more robust NHP HSPC engraftment and antibody production from liver and spleen. (A) MISTRG6 mice were xenotransplanted using the same protocol applied in MISTRG mice, this time allowing up to 72 hours from birth for HSPC injections. Necropsy was performed at an earlier timepoint to allow the detection of 10-1074-producing cells while antibody titers were still detectable. Control mice received unedited CD34^+^ cells. (B) NHP engraftment levels in the peripheral blood of MISTRG6 mice over time. (C) B cells (NHP-CD4^-^/CD8a^-^/CD20^+^) as a fraction of NHP lineage in the peripheral blood of MISTRG6 mice. (D) Anti-HIV antibody titers in the plasma of MISTRG6 mice as measured by ELISA. (E-G) 10-1074-expressing cells (Strep Tag II^+^), IgG-producing cells (NHP-IgG^+^/CD138^+^), and plasma cells (NHP-CD20^-/low^/CD31^+^/CD138^+^) in different tissues of MISTRG6 mice at necropsy. (H) Fraction of total 10-1074-expressing cells found across all tissues that are either B cells, antibody-producing cells, or plasma cells. (I) Editing levels at the target locus in peripheral blood. n = 8 mice, n = 3 NHP donors. Error bars represent SD. Red lines in (B-C) represent immunizations.

Engraftment levels in the non-viral condition were no different than control, whereas mice in the AAV6.2 template group displayed significantly lower NHP-CD45^+^ engraftment levels in the peripheral blood (Figure 5B). As anticipated, B cell populations remained high in MISTRG6 mice (Figure 5C). Anti-HIV antibody titers were detectable in five of seven mice at week 8, remained detectable in four of six mice at week 10, and were significantly above control mice receiving the same immunization regimen (Figure 5D). Plasma titers peaked at 4.0 µg/mL at 8 weeks post-transplantation. There was no significant difference in antibody titers between viral and non-viral conditions.

Cell surface expression levels of 10-1074 antibody remained very low in circulation, as did the number of NHP-IgG^+^/CD138^+^ antibody-producing cells (Figure S8). In contrast, 10-1074- and IgG-producing cells were present in the liver and spleen (Figure 5E-F). Plasma cells, as defined by CD20^low/-^/CD31^+^/CD138^+^ cells in NHP, were also detected in the bone marrow and liver (Figure 5G). Of note, one mouse died in each condition, including experimental and control groups, prior to the endpoint. Deceased mice showed clear signs of GVHD. Of all 10-1074-producing cells detected across tissues, 27-34% were CD20^+^ B cells and 5.3-13% were CD20^low/-^/CD31^+^/CD138^+^ plasma cells (Figure 5H).

Sequencing once again confirmed editing at the target NHP *IGH* locus in different tissues of transplanted mice, with AAV-edited HSPC displaying a higher proportion of large insertions (Figure 5I). Editing at the target locus averaged 13.7% in the peripheral blood of dsDNA mice and 18.4% in that of AAV mice, remaining constant over 2 weeks. Edited NHP cells were also detected in all tissues analyzed at necropsy (Figure S9). To confirm productive knock-in of the template in the non-viral condition, long-read sequencing of the *IGH* locus was performed on CD34^+^ cells. Amplicons of over 5.2 kb contained the full-length template at the expected insertion site, including an intact coding region, single-base changes unique to the homology arms, such as the blocking mutation at the PAM, and an additional 2.6 kb of the *M. nemestrina* genome past the homology arms, matching unedited samples (Sequence S1).

## DISCUSSION

This study lays the foundation to address a gap in current HIV therapies, which lack the ability to produce drugs or antibodies for extended periods of time and potentially lasting years. Existing genome engineering approaches for durable biologic production primarily rely on viral vectors. While effective, these methods are costly and can induce immune responses that limit practicality. By contrast, non-viral gene knock-in strategies that leverage endogenous antibody-producing cells and genomic loci could offer a promising alternative for producing engineered antibodies against HIV over a lifetime. Here, we utilized a relevant pre-clinical model of HIV to demonstrate that non-viral HSPC editing at the *IGH* locus offers a viable pathway to achieve expression of therapeutic antibodies.

Our data demonstrate that both CRISPR/Cas9 and CRISPR/Cas12a systems can generate genetic edits at the *IGH* locus in pigtail macaque primary HSPC. We observed that the choice of nuclease and gRNA influenced knock-in efficiency, extending beyond total editing efficiency. This finding aligns with previous studies suggesting that HDR outcomes may be influenced by the sequence context surrounding the cut site.^31,32^ Our data suggest that the staggered cuts produced by Cas12a, in contrast to the blunt double-strand breaks created by Cas9, may increase the likelihood of productive transgene knock-in or reduce off-target mutations at this locus. In addition to targeting a non-coding region, precise edits could minimize impacts on non-antibody producing cells derived from edited HSPC, improving the safety profile of this approach, but remain to be tested. We observed that the choice of template also impacted editing efficiency and productive transgene knock-in. Although it produced perfect HDR events with small templates (<200 bp), ssODN was less effective for achieving knock-in of large sequences (>200 bp) at the *IGH* locus. This is in contrast to other studies where knock-in was achieved in HSPC using ssODN.^38,39^ Though less efficient at gene knock-in at the human hemoglobin subunit beta (*HBB*) locus, ssODN templates did not impair engraftment as compared to viral templates with known genotoxic risks in humanized mouse models (NSG and NBSGW).^40^ It is of note that knock-in of anti-HIV bnAbs was recently reported in human HSPC with high efficiency at the *CCR5* locus using AAV following transplantation in NSG and NBSGW mice.^41^ Likely all of these factors together (choice of nuclease, target locus, DNA template, and mouse model) impact desired editing outcomes observed in primary cells. Our data in MISTRG/MISTRG6 mice suggest that HSPC edited *ex vivo* with long dsDNA templates at an endogenous antibody locus remain capable of engraftment without compromising productive knock-in efficiency and potential to differentiate into antibody-producing B cells.

The potential of gene-modified HSPC for treating chronic diseases is generating considerable interest, with authorization recently given for the first CRISPR-based gene therapy drug product including primary human HSPC for the treatment of sickle cell disease.^42^ However, the ability of CRISPR-mediated gene knock-ins in HSPC at highly recombinatory loci to persist throughout differentiation remains uncertain. Our results show that non-viral knock-in edited NHP HSPC are capable of engrafting into an immunodeficient mouse model, where they successfully differentiated into B cells capable of expressing antibody transgenes. Moreover, we detected antibody titers in the circulation of these mice, confirming that edited HSPC could differentiate into functional immune cells producing detectable levels of antibodies in response to antigen. While edited cells could express the transgene, their populations diminished over time, even with continued immunization. While these declines aligned with a well-described switch from peripheral B cells to T cells in humanized and primatized mice,^26,43^ it may also suggest selective pressure against edited cells, contributing to their reduction over time. It was recently reported that HSPC edited with Cas9 nuclease and ssODN template for HDR *ex vivo* demonstrated impaired long-term engraftment in a NHP model.^44^ This raises the question of whether dependence on the HDR pathway could impede clinical translation. One possible workaround could be the adoption of gene knock-in technologies that do not depend on endogenous repair machinery, such as systems based on engineered prime editors and recombinases.^45^

In initial experiments with MISTRG mice, the relatively low number of HSPC with large insertions transplanted, corresponding to approximately 1,000 cells (only 1 of which on average contains a perfect HDR event), likely failed to generate or spontaneously lost detectable bnAb levels due to the stochastic loss of HSPC clones during hematopoiesis. Although not a head-to-head comparison of the two strains, a change to MISTRG6 mice expressing human IL-6 was sufficient to increase the fraction of mice demonstrating plasma antibody titers from one of five mice to seven of seven mice. In addition, it has been shown that 10-1074 titers of 5 µg/mL, similar to those achieved here, are sufficient to reduce simian-HIV (SHIV) infection by ≥99.9% in NHP.^46^ Of note, we validated bnAb expression in HEK 293E; expression levels could be further optimized in mature primary NHP B cells. It also remains unclear how NHP antibody-producing B cells are maturing in the primatized MISTRG6 mouse model. There is limited evidence for the formation of germinal centers and long-lived plasma cells in humanized MISTRG6 mice.^47,48^ It remains unclear, however, whether optimization in mice to achieve higher numbers of antibody-producing animals is relevant to translation of this approach in the autologous setting in NHP. The use of better immunization regimens and the presence of fully functional germinal centers in an NHP model may produce biologically relevant antibody titers.

A non-viral approach to gene knock-in could have significant clinical value, particularly if adapted for *in vivo* delivery. HIV treatments such as ART can complicate viral transduction.^6^ A vector capable of delivering a RNP and DNA template to specific cell types for targeted, transient genome editing would offer a highly programmable platform for *in vivo* biologic production. In conclusion, this study provides foundational insights for the development of non-viral gene therapies aimed at achieving durable, *in vivo* antibody production by targeting HSPC. These findings could have broader implications for treating HIV and other chronic diseases through sustained biologic production, representing a significant advance in genome engineering and therapeutic development.

## MATERIALS AND METHODS

### HSPC isolation and culture

The NHP HSPC isolation protocol was derived from published methods and adapted for non-mobilized, healthy adult NHP donors.^49^ Bone marrow aspirates were obtained from the femur or humerus and hemolyzed (Pharm Lyse; BD Biosciences) the same day. White blood cells were enriched for CD34^+^ by sequentially incubating with 12.8 IgM anti-NHP-CD34 antibody (obtained from Fred Hutchinson Cancer Center Therapeutics Products Program,^50^) for 30 minutes at 4°C, followed by anti-IgM microbeads (Miltenyi Biotec) for 30 minutes at 4°C, and then loading onto a magnetic column (Miltenyi Biotec). NHP-CD34^+^ cells were eluted from the column and prepared for flow cytometry to confirm purity (Table S2).

Immediately following isolation, NHP HSPC were transferred to HSPC medium (StemSpan Serum-Free Expansion Medium (SFEM) II (StemCell Technologies) with 100 ng/mL recombinant human thrombopoietin (TPO), stem cell factor (SCF), and flightless 3 (FLT-3) ligand, all from CellGenix) at 1.0×10^6^ cells/mL in tissue culture (TC)-treated flasks, incubated at 37°C and 5% CO_2_. Alternatively, NHP HSPC were cryopreserved in CS10 medium (StemCell Technologies) or in 90% heat-inactivated fetal bovine serum (FBS; Gibco) with 10% dimethyl sulfoxide (DMSO; Thermo Fisher Scientific).

### *IGH* sequencing and gRNA selection

Sequencing primers for NHP *IGH* were designed based on the rhesus macaque genome (rheMac10 assembly, Table S3). Genomic DNA was isolated from pigtail macaque and used to amplify 1,024 bp segments, which were submitted for Sanger sequencing. Pigtail macaque sequences were aligned using MAFFT (v7)^51^ to identify divergence from rhesus macaque and human sequences.

All possible CRISPR/Cas9 and CRISPR/Cas12a gRNA within this sequence were computationally screened using CRISPOR (v5.01).^33^ These gRNA were ranked by predicted specificity scores, efficiency scores, and Lindel scores (logistic regression model to predict insertions and deletions) when available, in that order (Table S1). The top 5 gRNA for each nuclease with guide and protospacer-adjacent motif (PAM) sequences that had perfect homology to both rhesus and pigtail macaques were selected for synthesis. CRISPR/Cas9 were synthesized as CRISPR RNA (crRNA) for pairing with tracer RNA (trRNA), while CRISPR/Cas12a were synthesized as single gRNA (Integrated DNA Technologies (IDT)).

### NHP B cell culture

B lymphocyte cell line (LCL) 8664 (American Type Culture Collection (ATCC)), a lymphoma-derived rhesus macaque cell line,^52^ were maintained in Roswell Park Memorial Institute (RPMI) medium 1640 containing 10% FBS and 1% penicillin/streptomycin (Gibco) at 37°C and 5% CO_2_. Cells were cultured in TC-treated flasks and passaged at 80-90% confluence approximately once every 4 days. Cell identity was authenticated by Sanger sequencing.

### HDR template construction

Templates for measuring HDR efficiency in NHP B cells were synthesized as ssODN (IDT) of the non-template strand with 40-bp homology arms flanking an 8-bp insertion. Templates for each gRNA were designed for insertion at the predicted cut site, namely −9 bp from Cas9 PAM or +15 bp from Cas12a PAM. Homology arms contained 2-bp blocking mutations in the PAM sequence for Cas9 gRNA and 1-bp blocking mutations for Cas12a gRNA to prevent cutting following HDR and multiple insertion events.

Templates for gene knock-in were cloned as bacterial plasmids containing ampicillin resistance. The 450-bp homology arms were amplified from pigtail macaque genomic DNA. As VH4 is the major expressed VH allele in rhesus macaque, alleles from the IgVH4 locus were aligned, the most conserved of which (VH4-38-2 allele) was selected for B cell-specific expression.^36^ Expression was confirmed in NHP B cells. Transgene sequences, including linker and splice signals, and a compact CMV promoter were synthesized as gBlocks (IDT).^53^ Cloning was performed by DNA ligation (Thermo Fisher Scientific), blocking mutations were introduced by site-directed mutagenesis, and all final products were confirmed by whole plasmid sequencing (Plasmidsaurus). Plasmids were used to PCR amplify gel-purified dsDNA templates and to synthesize long ssODN templates (GenScript). All templates will be made available through Addgene upon publication.

### RNP electroporation

LCL 8664 cells were passaged to obtain a cell population of 1.0 × 10^6^ cells per sample in the exponential growth phase on the day of electroporation. Cas9 RNP complex was prepared by combining 3 µL 200 µM crRNA, 3 µL 200 µM tracrRNA, and 6 µL of duplex buffer (IDT). The mixture was incubated at 95°C for 5 minutes, followed by cooling at room temperature for 15 minutes. Cas12a RNP complex was prepared by combining 3 µL 200 µM crRNA, 3 µL water, and 6 µL of duplex buffer. To each reaction, 4 µL 100 µM DNA template and 1 µL 10 mg/mL Cas9 or Cas12a nuclease (Aldevron) were added and incubated at room temperature for 5 minutes. For the control group, templates were electroporated without nuclease. Cells were recovered in culture for 3 days prior to genomic DNA isolation for sequencing analysis.

Cryopreserved primary NHP HSPC were thawed in Iscove’s Modified Dulbecco’s Medium (IMDM), recovered for 24 hours in HSPC medium at 1.0 ×10^6^ cells/mL in a TC-treated T25 flask, and washed twice with Dulbecco’s phosphate-buffered saline (PBS; Gibco) prior to electroporation. For gRNA screening, HSPC were electroporated using an ECM 830 Electroporation System (Harvard Bioscience). In each reaction, 5.0 × 10^5^ CD34^+^ cells were combined with 10 nmol gRNA and 180 pmol nuclease in 50 µL BTX buffer (Harvard Bioscience), and electroporated at 125 V for 5 ms. Cells were recovered in 1.0 mL HSPC medium in a TC-treated 12-well plate for 3 days, after which genomic DNA was isolated for sequencing.

For the MISTRG mice receiving reporter edited cells, HSPC were electroporated using an Amaxa Nucleofector II Device (Lonza). In each reaction, 5.0 × 10^5^ CD34^+^ cells were combined with 10 nmol gRNA, 180 pmol nuclease, and 5 µg dsDNA template in 100 µL Nucleofector Solution V with Supplement 1 from Nucleofector Kit V (Lonza), and electroporated using program U-008. For MISTRG mice receiving antibody edited cells, HSPC were electroporated using a Neon Transfection System (Thermo Fisher Scientific). In each reaction, 5.0 × 10^5^ CD34^+^ cells were combined with 10 nmol gRNA, 180 pmol nuclease, and 5 µg dsDNA template or 5.0 × 10^11^ viral genomes (vg) AAV6.2 template in 100 µL buffer T (Thermo Fisher Scientific), and electroporated in a 100 µL tip at 1650 V for 3 pulses with a width of 10 ms. In both cases, cells were recovered in 1.5 mL HSPC medium in a TC-treated 6-well plate for 4 hours prior to injection or for 3 days prior to genomic DNA isolation for sequencing.

### Indel and HDR analysis

Genomic DNA samples were prepared for high-throughput sequencing on a MiSeq platform (Illumina). PCR amplifications were performed to enrich the NHP *IGH* locus using target-specific primers (Table S3), to then attach overhangs using P5 and P7 adapters, and finally to barcode each sample using Nextera indexes (Illumina), gel purifying amplicons of the desired length at each step. Indexed amplicons were quantified and pooled at 5 or 10 nM. Paired-end, 250-bp reads were sequenced using MiSeq Nano v2 according to the manufacturer’s protocol.

Sequencing results were processed using an in-house algorithm and adapted to the target NHP reference sequence (GitHub: https://github.com/jack-cast/FredHutch_Gene_Edit_2).^54^ Stitched reads are aligned to the reference sequence. Indels and HDR events are measured within a quantification window spanning 50-bp on either side of the gRNA and normalized to a control, non-edited sample. Due to heterogeneity at the *IGH* locus in LCL 8664 cells, indel and HDR percentages were calculated per target allele with perfect sequence identity for each gRNA.

### AAV transduction

Based on previous literature, 4 serotypes were chosen for testing on NHP-CD34^+^ cells: AVV2, AAV6, AAV6.2, and AAV-DJ.^55^ A standard *in vitro* grade panel with CMV-mediated GFP reporter expression (VectorBuilder) was transduced into 3.75 × 10^5^ cells at multiplicities of infection (MOI) of 2.4 × 10^4^ and 6.7 × 10^5^ vector particles (vp). Cell viability and GFP expression were assessed by flow cytometry at days 1 and 3 post-transduction. Based on these results, the AAV6.2 serotype was selected for packaging of the VH4-10-1074^Cas12a^ template. For HSPC transduction prior to mouse transplantation, AAV6.2 was added at an MOI of 1.0 × 10^6^ vp alongside RNP during electroporation.

### Primatized mouse studies

For xenotransplantation experiments, cryopreserved NHP HSPC were slow thawed within 24 hours of MISTRG/MISTRG6 mouse birth and recovered in HSPC medium overnight. Within 48 hours (MISTRG) or 72 hours (MISTRG6) of birth and 4 hours prior to injection, pups are irradiated with 150 cGy γ-rays in a cesium-137 irradiator. NHP HSPC from different donors were pooled together, electroporated as described above, recovered for 4 hours in HSPC media, then resuspended in PBS with 1% heparin (Table S4). HSPC were injected intra-hepatically at variable cell doses in 40 µL. Control HSPC were transferred to the same 6-well plate without electroporation. When cell counts allowed, non-injected HSPC samples were set aside for 3-day recovery and genomic DNA isolation.

Peripheral blood from the retro-orbital sinus at various time points was collected into Vacutainer K2 EDTA coated tubes (BD Biosciences). When applicable, plasma samples were collected by centrifuging at 1,000 x g for 10 minutes and transferring the supernatant either to be stored at 4°C or in aliquots at −20°C. Without plasma collection, blood samples were centrifuged at 600-800 g for 3 minutes. Cell pellets were treated twice with 1.0 mL of ACK Lysis Buffer (Thermo Fisher Scientific), first for 5 minutes then for 3 minutes. Cells were washed and resuspended in 1.0 mL of PBS. Approximately half of the sample was pelleted and reserved for genomic DNA analysis, the other half of the sample was further processed for flow cytometry staining. During terminal cardiac puncture bleeds, blood was collected in test tubes and allowed to clot for 30 minutes. Serum from the supernatant was then stored at 4°C.

Liver, spleen, femurs, and, when available, thymus and lymph nodes of mice were harvested at necropsy. Livers and spleens were processed into single cell suspensions by grinding the tissue between the frosted edges of two microscope slides. Bone marrow was isolated from femurs by removing all excess tissue around the bone and snipping at the trochanter and epicondyle ends. A 27-gauge needle was inserted through the medullary cavity, and the cavity was flushed with 1.0 mL of PBS twice. Thymus and lymph nodes were processed into single cell suspensions by using the back end of the plunger of a 3-mL syringe to pass the tissue through a 100 µm cell strainer placed in a dish with PBS. Cell suspensions were filtered through a 100 µm cell strainer and washed with PBS. Of each sample type, approximately 2.0 × 10^5^ cells were reserved for flow cytometry, 2.0 × 10^5^ cells were processed for genomic DNA extraction, and 2.0 × 10^5^ cells were cryopreserved in 90% FBS and 10% DMSO.

### Recombinant bnAb production

The transgene for 10-1074 bnAb was cloned into a mammalian expression plasmid (pTT3) containing a NHP IgG constant heavy chain with a 6x His-Avi-Tag using In-Fusion (Takara Bio^56^). The resulting plasmid (pTT3 10-1074 NHP-IgG) expressed full length antibody and was confirmed by whole plasmid sequencing (Plasmidsaurus).

Human embryonic kidney (HEK) 293E cells, which are engineered for enhanced protein expression, were maintained in RPMI media containing 10% FBS and 1% penicillin/streptomycin at 37°C and 5% CO_2_.^56^ One day prior to transfection, 1.6 × 10^7^ cells were seeded in a TC-treated T150 flask with Freestyle 293 Expression Medium (Gibco) at a density of 5.0 × 10^7^ cells/mL. 10 µg of pTT3 10-1074 NHP-IgG was transfected using 40 µL 293-Free Transfection Reagent (MilliporeSigma). Cell cultures were grown out for 7 days, after which recombinant protein was isolated from the supernatant using Protein A IgG Purification Kit (Thermo Fisher Scientific).

### Bio-layer interferometry

HIV gp120 monomers from various clade C strains – including 16055-2 (accession ABL67444.1, Abcam), Du172.17 (accession DQ411853), ZA.1197MB (accession AY463234), CN97001 (accession G4XFJ5-1 with E46G, T396A, A497T), and ZM249M.PL1 (accession DQ388514, Thermo Fisher) – were screened for kinetic interactions with recombinant 10-1074 using the Octet RED96 Instrument (Sartorius). Anti-Human Fc Capture (AHC) Biosensors (Sartorius) were soaked for 15 minutes in Kinetics Buffer consisting of 1% Bovine Serum Albumin (BSA; Thermo Fisher Scientific), 0.05% sodium azide (MilliporeSigma), and 0.02% polysorbate 20 (Tween20; MilliporeSigma) in PBS. 1 µM of antigen and 40 µg/mL of antibody were loaded onto a non-treated, clear bottom 96-well plate. A non-specific antibody (MxR01, courtesy of Matthew Gray) was used as a negative control. AHC probes were loaded onto the instrument and allowed to equilibrate in the kinetics buffer. Once confirmed, 10-1074 was loaded onto the probe, and a baseline with the loaded antibody was confirmed. The association of the antibody to various antigens was then measured. To determine dissociation, probes were dipped into analyte free kinetics buffer. Association Rate Constant (K_a_), Dissociation Rate Constant (K_d_), and Affinity Constant (K_D_) were calculated, and the control antibody sample was adjusted to be the baseline standard for all antigens using the Octet Software (Sartorius, version 3.43).

### Flow cytometry

Flow cytometry samples were first stained for dead cells by incubating in Zombie-NIR Live/Dead Stain (BioLegend) diluted 1:250 in PBS for 15-20 minutes. Cells were washed with 1.0 mL FACS buffer (1mM EDTA (Thermo Fisher Scientific) and 0.5% FBS in PBS), and mouse samples were incubated in Fc block (anti-mouse CD16/CD32, clone 2.4G2, BD Biosciences) diluted 1:25 in FACS buffer for 10 minutes. Antibodies used in the various flow panels were combined in FACS buffer and added to cells for 20-30 minutes (Tables S5-6). Samples were washed and resuspended in FACS buffer and run on a FACSCelesta for cell cultures or a FACSymphony A5 for mouse samples (BD Biosciences). Results were analyzed to exclude dead cells and debris, and to sort live cells by a standard gating strategy using FlowJo (TreeStar, version 10.8.0, Figures S10-11).

### Kinetic colorimetric ELISA

Nunc immuno clear bottom 96-well plates (Thermo Fisher Scientific) were coated with 100 ng per well of HIV gp120 (426c Core with biotin, courtesy of Matthew Gray) in 100 µL PBS. Freshly coated plates were sealed and shaken at 250 rpm for 5-10 minutes to evenly distribute antigen and then incubated overnight at 4°C. Post-incubation, all wells were washed three times with wash buffer (0.05% Tween20 in PBS). Plates were blocked by incubating in 150 µL blocking buffer (wash buffer with 3.0% BSA) and shaking at 250 rpm for 1 hour at room temperature. Excess blocking buffer was removed and 100 µL of recombinant 10-1074 was loaded onto the plate in serial dilutions ranging in concentrations from 1-2,048 ng/mL as standards. Plasma samples were loaded onto the plate at a 1:50 dilution in blocking buffer. The plate was shaken at 250 rpm for 90 minutes at room temperature. The plate was washed 3 times with wash buffer, and goat anti-rhesus IgG(H+L)-horseradish peroxidase (HRP, SouthernBiotech) at a 1:6,000 dilution (MISTRG plasma) or a 1:3,000 dilution (MISTRG6 plasma) in blocking buffer was added to the wells and incubated for 1 hour. The plate was washed 3 times with wash buffer, and 100 µL of 3,3’,5,5’-tetramethylbenzidine (TMB) solution (Thermo Fisher Scientific) was added to each well. Samples and standards were then analyzed at 2.5-minute increments up to 30 minutes using a BioTek Synergy H4 Hybrid Multi-Mode Microplate Reader (Marshall Scientific) set at a 405 nm absorbance wavelength.

### Data Availability

Data are included within the paper and its supplemental information. Raw data collected are available from the corresponding author upon request.

## SUPPLEMENTAL INFORMATION

Tables S1-S6.

Figures S1-S11.

Sequence S1.

## ACKNOWLEDGMENTS

The authors would like to thank Audrey Germond and William D. Garrison for support in obtaining NHP materials, Ekram Gad for assisting with mouse studies, Jonathan Linton and Cyd N. McKay for providing MISTRG mice, and Rachel A. Bender Ignacio and Helen Crawford for reviewing the manuscript. Research reported in this publication was supported by the National Institutes of Health Office of the Director, Office of Research Infrastructure Programs (ORIP) under award number P51OD010425 and U42OD011123. The content is solely the responsibility of the authors and does not necessarily represent the official views of the National Institutes of Health. J. Adair is The Fleischauer Family Endowed Chair in Gene Therapy Translation. This research was also supported by the Genomics and Bioinformatics Shared Resource (RRID:SCR_022606), the Comparative Medicine Shared Resource (RRID:SCR_022610), and the Flow Cytometry Shared Resource (RRID:SCR_022613) of the Fred Hutch/University of Washington/Seattle Children’s Cancer Consortium (P30 CA015704). This study was supported by NIH grant R01AI167009 (MPIs: J. Adair and J. Taylor), and by a grant from the Bill and Melinda Gates Foundation (INV002613; PI J. Adair).

## AUTHOR CONTRIBUTIONS

J.M.P.C, J.J.T., and J.E.A. conceived ideas. J.M.P.C. and J.E.A. designed research. J.M.P.C. and K.P. wrote the paper. J.E.A. and J.J.T. supervised this research, secured funding, and edited the paper. J.M.P.C, K.P., Y.J., M.E.C., M.D.G., and J.N.S.G. performed research. M.D.G. produced and contributed plasmids and proteins for this study. J.M.P.C. and M.R.E. performed sequencing analysis. J.D.L. and A.R. bred and contributed mice and input into murine studies. All authors read and approved the manuscript.

