## Supplemental Information for "*In vivo* production of an anti-HIV antibody from primate hematopoietic cells by non-viral knock-in"

### LIST OF SUPPLEMENTAL INFORMATION

Tables S1-S6.

Figures S1-S11.

Sequence S1.

### 13 SUPPLEMENTARY TABLES

**Table S1. Cas9 and Cas12a gRNA sequences and scores.**

| Sequence (including PAM) | Nuclease | Specificity score (MIT) | Efficiency score (Doench '16 for Cas9, DeepCpf1 for Cas12a) | Lindel score | PM | RM | Synthesized as |
|---|---|---|---|---|---|---|---|
| TTTTTTTAACTGTCTAAGTATTTG | Cas12a | 75.5 | 1.4 | | TRUE | | |
| TTTGGATTCCTGGGGCCAAGGGGC | Cas12a | 64.0 | 51.4 | | TRUE | | |
| TTTTTTAACTGTCTAAGTATTTGA | Cas12a | 57.7 | 1.4 | | TRUE | | |
| TTTTTAACTGTCTAAGTATTTGAA | Cas12a | 56.5 | 9.7 | | TRUE | | |
| GACATGTTCCAAGGGCACCTGGG | Cas9 | 56.4 | 62.4 | 88.0 | TRUE | | |
| TTTAACTGTCTAAGTATTTGAAAT | Cas12a | 55.6 | 9.7 | | TRUE | | |
| TTTTAACTGTCTAAGTATTTGAAA | Cas12a | 55.1 | 28.4 | | TRUE | | |
| TTTCAAATACTTAGACAGTTAAAA | Cas12a | 54.6 | 46.5 | | TRUE | | |
| AGAAAACAAAGGCTCTAGAGTGG | Cas9 | 52.8 | 61.4 | 81.0 | TRUE | TRUE | NHP_IGH_Cas9_1 |
| CCAGTCCGCCCAGGTGCCCTTGG | Cas9 | 51.6 | 37.6 | 68.0 | TRUE | | |
| TTTCTGGACTACCCGTGCCCCGAC | Cas12a | 49.9 | 29.8 | | TRUE | TRUE | NHP_IGH_Cas12a_1 |
| TTTGGACGAGATGCCAGAGCAAAC | Cas12a | 49.7 | 73.9 | | TRUE | TRUE | NHP_IGH_Cas12a_2 |
| TTTAGAATTATGAGGTGCGCTGTG | Cas12a | 49.5 | 30.5 | | TRUE | TRUE | NHP_IGH_Cas12a_3 |
| TTTTGAAGTATATTAATTTTTTTA | Cas12a | 49.2 | 1.1 | | TRUE | | |
| TTTAATAAGCACGCCTCTTAAGAT | Cas12a | 49.2 | 33.5 | | TRUE | TRUE | NHP_IGH_Cas12a_4 |
| TTTCTTTAGAATTATGAGGTGCGC | Cas12a | 49.1 | 2.1 | | TRUE | TRUE | NHP_IGH_Cas12a_5 |
| TTTGCTCTGGCATCTCGTCCAAAT | Cas12a | 49.0 | 51.5 | | TRUE | TRUE | |
| TTTGTGGGGTGAGGATGGACATTC | Cas12a | 48.9 | 31.8 | | TRUE | TRUE | |
| TTTTGTGGGGTGAGGATGGACATT | Cas12a | 48.9 | 55.0 | | TRUE | | |
| TTTAACATTTAATAAGCACGCCTC | Cas12a | 48.8 | 14.5 | | TRUE | TRUE | |
| TTTGATTAACACCCATGAGTGGTA | Cas12a | 48.7 | 34.4 | | TRUE | TRUE | |
| TTTTCCAAAGGCATCGGAAAATCC | Cas12a | 48.7 | 54.7 | | TRUE | TRUE | |
| TTTGGAAAATGGGACTTAGGTTGG | Cas12a | 48.7 | 57.1 | | TRUE | TRUE | |
| TTTTCTGAGCATTGCAGGCTGGTC | Cas12a | 48.6 | 36.8 | | TRUE | TRUE | |
| TTTTCTGCTATTGCCTGTGGGGTT | Cas12a | 48.6 | 27.6 | | TRUE | TRUE | |

**Table S1. Cas9 and Cas12a gRNA sequences and scores.**

| Sequence (including PAM) | Nuclease | Specificity score (MIT) | Efficiency score (Doench '16 for Cas9, DeepCpf1 for Cas12a) | Lindel score | PM | RM | Synthesized as |
|---|---|---|---|---|---|---|---|
| TTTTCTTTAGAATTATGAGGTGCG | Cas12a | 48.6 | 2.4 | | TRUE | TRUE | |
| TTTCTGAGCATTGCAGGCTGGTCC | Cas12a | 48.5 | 16.7 | | TRUE | TRUE | |
| TTTTTGTGGGGTGAGGATGGACAT | Cas12a | 48.5 | 31.8 | | TRUE | | |
| TTTCATGATTCGCTGTGTGTGCAG | Cas12a | 48.5 | 27.6 | | TRUE | TRUE | |
| GGACATGTTCCAAGGGCACCTGG | Cas9 | 48.4 | 37.9 | 76.0 | TRUE | | |
| TTTCCGATGCCTTTGGAAAATGGG | Cas12a | 48.4 | 43.2 | | TRUE | TRUE | |
| TTTTCTCTGGAGCCACTTCAAACA | Cas12a | 48.4 | 46.0 | | TRUE | TRUE | |
| TTTGGCCCTAATTCCAGAGACATA | Cas12a | 48.3 | 78.7 | | TRUE | TRUE | |
| CCAGGACGGATGCGTAGCCTTGG | Cas9 | 48.2 | 49.4 | 78.0 | TRUE | TRUE | NHP_IGH_Cas9_2 |
| TTTCTCTGGAGCCACTTCAAACAT | Cas12a | 48.1 | 28.5 | | TRUE | TRUE | |
| TTTCTCTTGGAAACCAACTTCAGG | Cas12a | 48.1 | 21.0 | | TRUE | TRUE | |
| TTTTTCTCTGGAGCCACTTCAAAC | Cas12a | 48.1 | 28.5 | | TRUE | TRUE | |
| TTTAAGATGCAGGTTGGCACACAG | Cas12a | 48.1 | 45.0 | | TRUE | TRUE | |
| TTTTAACATTTAATAAGCACGCCT | Cas12a | 48.0 | 19.4 | | TRUE | TRUE | |
| GGCTACGCATCCGTCCTGGCTGG | Cas9 | 47.9 | 46.4 | 77.0 | TRUE | TRUE | NHP_IGH_Cas9_3 |
| TTTTCCGATGCCTTTGGAAAATGG | Cas12a | 47.9 | 48.0 | | TRUE | TRUE | |
| TTTCCAAAGGCATCGGAAAATCCA | Cas12a | 47.9 | 39.9 | | TRUE | TRUE | |
| CCAAGGCTACGCATCCGTCCTGG | Cas9 | 47.7 | 52.2 | 85.0 | TRUE | TRUE | NHP_IGH_Cas9_4 |
| TTTCTGCTATTGCCTGTGGGGTTT | Cas12a | 47.6 | 4.7 | | TRUE | TRUE | |
| TTTGTTGGCTGGAAAGAGAACTGT | Cas12a | 47.2 | 19.4 | | TRUE | TRUE | |
| TTTTAAGTCATAATTGTCTTAACC | Cas12a | 47.0 | 41.0 | | TRUE | TRUE | |
| TTTTCTCTTGGAAACCAACTTCAG | Cas12a | 46.9 | 35.3 | | TRUE | TRUE | |
| TTTAAGTCATAATTGTCTTAACCA | Cas12a | 46.4 | 11.5 | | TRUE | TRUE | |
| TGCCCTTGGAACATGTCCCGAGG | Cas9 | 46.3 | 64.0 | 85.0 | TRUE | | |
| TTTGTTTTCTGCTATTGCCTGTGG | Cas12a | 46.2 | 2.5 | | TRUE | TRUE | |
| TTTGAAGTGGCTCCAGAGAAAAAT | Cas12a | 46.1 | 60.1 | | TRUE | TRUE | |
| TTTCCAGCCAACAAAGAATTTAAG | Cas12a | 46.0 | 60.9 | | TRUE | TRUE | |
| TTTTTTAACTGTCCAAGTATTTGA | Cas12a | 45.9 | 1.5 | | | TRUE | |

**Table S1. Cas9 and Cas12a gRNA sequences and scores.**

| Sequence (including PAM) | Nuclease | Specificity score (MIT) | Efficiency score (Doench '16 for Cas9, DeepCpf1 for Cas12a) | Lindel score | PM | RM | Synthesized as |
|---|---|---|---|---|---|---|---|
| ACCCATGAGTGGTATGTCTCTGG | Cas9 | 45.7 | 41.4 | 81.0 | TRUE | TRUE | NHP_IGH_Cas9_5 |
| TTTTTAAGTCATAATTGTCTTAAC | Cas12a | 45.4 | 11.5 | | TRUE | TRUE | |
| TTTCCAAGAGAAAAGGATTGTTCA | Cas12a | 44.8 | 57.1 | | TRUE | TRUE | |
| GTCCTCGGGACATGTTCCAAGGG | Cas9 | 44.8 | 62.1 | 71.0 | TRUE | TRUE | |
| TCCCCTTGGAACATGTCCCGAGG | Cas9 | 44.7 | 65.2 | 86.0 | | TRUE | |
| TTTTTAACTGTCCAAGTATTTGAA | Cas12a | 44.2 | 14.4 | | | TRUE | |
| ATGTTCCAAGGGCACCTGGGCGG | Cas9 | 44.1 | 69.3 | 82.0 | TRUE | | |
| TCCTCGGGACATGTTCCAAGGGG | Cas9 | 43.8 | 65.8 | 78.0 | | TRUE | |
| TTTAACTGTCCAAGTATTTGAAAT | Cas12a | 43.8 | 14.4 | | | TRUE | |
| GGGGCACGGGTAGTCCAGAAAGG | Cas9 | 43.7 | 54.8 | 83.0 | TRUE | TRUE | |
| GTCTCAGGTGCGGTGTCTGTAGG | Cas9 | 43.5 | 55.8 | 78.0 | TRUE | TRUE | |
| CCAAGGGGACCTGGGCGGACTGG | Cas9 | 43.5 | 40.5 | 53.0 | | TRUE | |
| CCAAGGGCACCTGGGCGGACTGG | Cas9 | 43.4 | 44.8 | 55.0 | TRUE | | |
| CTAAGACCCCTGGTTTGCTCTGG | Cas9 | 43.2 | 47.0 | 87.0 | TRUE | TRUE | |
| GGTCCTCGGGACATGTTCCAAGG | Cas9 | 42.8 | 52.2 | 73.0 | TRUE | TRUE | |
| TCCAGAGACATACCACTCATGGG | Cas9 | 42.8 | 59.7 | 84.0 | TRUE | TRUE | |
| CAAACCAGGGGTCTTAGTGATGG | Cas9 | 42.8 | 54.6 | 76.0 | TRUE | TRUE | |
| TTTCAAATACTTGGACAGTTAAAA | Cas12a | 42.5 | 47.4 | | | TRUE | |
| TGTGGATTTTCCGATGCCTTTGG | Cas9 | 42.3 | 33.4 | 70.0 | TRUE | TRUE | |
| TTTTAACTGTCCAAGTATTTGAAA | Cas12a | 42.2 | 31.2 | | | TRUE | |
| TCTTGATGAGAGCAGGGTCGGGG | Cas9 | 42.2 | 53.5 | 76.0 | TRUE | TRUE | |
| TTTCACATTTTTAAGTCATAATTG | Cas12a | 42.0 | 7.5 | | TRUE | TRUE | |
| CCTTCCTGGCCAGTCCGCCCAGG | Cas9 | 41.9 | 45.8 | 70.0 | TRUE | TRUE | |
| GGGCACGGGTAGTCCAGAAAGGG | Cas9 | 41.6 | 49.7 | 75.0 | TRUE | TRUE | |
| TCTGGCATCTCGTCCAAATGTGG | Cas9 | 41.6 | 61.9 | 82.0 | TRUE | TRUE | |
| TTTGAAATTCTTATCATTTGATTA | Cas12a | 41.1 | 42.0 | | TRUE | TRUE | |
| TTTTGAAGTATATTAACTTTTTTA | Cas12a | 40.9 | 3.1 | | | TRUE | |
| TTCCAGAGACATACCACTCATGG | Cas9 | 40.9 | 54.7 | 80.0 | TRUE | TRUE | |

**Table S1. Cas9 and Cas12a gRNA sequences and scores.**

| Sequence (including PAM) | Nuclease | Specificity score (MIT) | Efficiency score (Doench '16 for Cas9, DeepCpf1 for Cas12a) | Lindel score | PM | RM | Synthesized as |
|---|---|---|---|---|---|---|---|
| GGCACCTGGGCGGACTGGCCAGG | Cas9 | 40.7 | 42.3 | 81.0 | TRUE | | |
| GCCTTGGTCTTGATGAGAGCAGG | Cas9 | 40.7 | 45.2 | 84.0 | TRUE | TRUE | |
| ATTTGATTAACACCCATGAGTGG | Cas9 | 40.6 | 64.1 | 85.0 | TRUE | TRUE | |
| TCAGCCATCACTAAGACCCCTGG | Cas9 | 40.4 | 56.0 | 79.0 | TRUE | TRUE | |
| GGACATGTTCCAAGGGGACCTGG | Cas9 | 40.4 | 34.6 | 68.0 | | TRUE | |
| ACGAGATGCCAGAGCAAACCAGG | Cas9 | 40.3 | 48.1 | 87.0 | TRUE | TRUE | |
| CTTAGGTTGGATGTGTCTGATGG | Cas9 | 39.7 | 54.3 | 74.0 | TRUE | TRUE | |
| GTTTTCTGAGCATTGCAGGCTGG | Cas9 | 39.6 | 48.6 | 85.0 | TRUE | TRUE | |
| GACGGGCACTGGGGTGCCTTGGG | Cas9 | 39.6 | 32.1 | 66.0 | TRUE | TRUE | |
| TGGGGTTTTCTGAGCATTGCAGG | Cas9 | 39.5 | 23.6 | 84.0 | TRUE | TRUE | |
| CAGAGTGGGTGAATCCAGCCAGG | Cas9 | 39.5 | 58.0 | 79.0 | TRUE | TRUE | |
| CACGGGTAGTCCAGAAAGGGTGG | Cas9 | 39.4 | 60.5 | 78.0 | TRUE | TRUE | |
| GGGACCTGGGCGGACTGGCCAGG | Cas9 | 39.4 | 41.8 | 81.0 | | TRUE | |
| GCTGTCCTGACAGAGGCTTAGGG | Cas9 | 39.4 | 58.5 | 80.0 | TRUE | TRUE | |
| CGAGATGCCAGAGCAAACCAGGG | Cas9 | 39.2 | 73.3 | 85.0 | TRUE | TRUE | |
| AGCATTGCAGGCTGGTCCTCGGG | Cas9 | 39.1 | 47.7 | 62.0 | TRUE | TRUE | |
| TTTGAAGTATATTAACTTTTTTAA | Cas12a | 39.0 | 21.8 | | | TRUE | |
| GGGGCTTGGGGAGCCACATTTGG | Cas9 | 38.9 | 25.5 | 70.0 | TRUE | TRUE | |
| TTCTCTTGGAAACCAACTTCAGG | Cas9 | 38.7 | 25.9 | 69.0 | TRUE | TRUE | |
| CCTTGGTCTTGATGAGAGCAGGG | Cas9 | 38.4 | 62.4 | 88.0 | TRUE | TRUE | |
| ACGGGCACTGGGGTGCCTTGGGG | Cas9 | 38.4 | 44.8 | 73.0 | TRUE | TRUE | |
| GGCCAGGAAGGGACGGGCACTGG | Cas9 | 38.2 | 35.7 | 72.0 | TRUE | TRUE | |
| GCCAGGAAGGGACGGGCACTGGG | Cas9 | 38.0 | 38.4 | 62.0 | TRUE | TRUE | |
| CTGGGCGGACTGGCCAGGAAGGG | Cas9 | 37.7 | 53.0 | 74.0 | TRUE | TRUE | |
| TTGGAAAATGGGACTTAGGTTGG | Cas9 | 37.7 | 39.7 | 80.0 | TRUE | TRUE | |
| GTCCTGACAGAGGCTTAGGGAGG | Cas9 | 37.6 | 61.7 | 76.0 | TRUE | TRUE | |
| CCAGGAAGGGACGGGCACTGGGG | Cas9 | 37.4 | 53.4 | 81.0 | TRUE | TRUE | |
| ATGTGGCTCCCCAAGCCCCCAGG | Cas9 | 37.3 | 46.7 | 81.0 | TRUE | TRUE | |

| Table S1. Cas9 and Cas12a gRNA sequences and scores. | | | | | | | |
|---|---|---|---|---|---|---|---|
| **Sequence (including PAM)** | **Nuclease** | **Specificity score (MIT)** | **Efficiency score (Doench '16 for Cas9, DeepCpf1 for Cas12a)** | **Lindel score** | **PM** | **RM** | **Synthesized as** |
| GAGCATTGCAGGCTGGTCCTCGG | Cas9 | 37.3 | 40.1 | 80.0 | TRUE | TRUE | |
| GGCCTCCCTAAGCCTCTGTCAGG | Cas9 | 37.2 | 50.7 | 83.0 | TRUE | TRUE | |
| ATGAGAGCAGGGTCGGGGCACGG | Cas9 | 37.2 | 43.5 | 77.0 | TRUE | TRUE | |
| TGAGGAATGTGTCTCAGGTGCGG | Cas9 | 37.1 | 70.7 | 69.0 | TRUE | TRUE | |
| GTCTTGATGAGAGCAGGGTCGGG | Cas9 | 37.1 | 45.4 | 85.0 | TRUE | TRUE | |
| TTTGAAGTATATTAATTTTTTTAA | Cas12a | 37.0 | 2.3 | | TRUE | | |
| ATCTTAAATTCTTTGTTGGCTGG | Cas9 | 37.0 | 44.1 | 81.0 | TRUE | TRUE | |
| AGGGGTCTTAGTGATGGCTGAGG | Cas9 | 37.0 | 62.1 | 80.0 | TRUE | TRUE | |
| AAAGGATTGTTCATCTTAAGAGG | Cas9 | 37.0 | 50.0 | 78.0 | TRUE | TRUE | |
| GTTTTCTGCTATTGCCTGTGGGG | Cas9 | 36.8 | 54.5 | 90.0 | TRUE | TRUE | |
| TAACTGAGCCTGGGGGCTTGGGG | Cas9 | 36.7 | 41.7 | 81.0 | TRUE | TRUE | |
| GTAACTGAGCCTGGGGGCTTGGG | Cas9 | 36.7 | 21.3 | 80.0 | TRUE | TRUE | |
| ACTGGGGTGCCTTGGGGATCTGG | Cas9 | 36.5 | 14.0 | 74.0 | TRUE | TRUE | |
| AGGACAGCAGCCACCCTTTCTGG | Cas9 | 36.3 | 19.7 | 75.0 | TRUE | TRUE | |
| TCTGATGGAGTAACTGAGCCTGG | Cas9 | 36.3 | 41.8 | 74.0 | TRUE | TRUE | |
| GAGATGCCAGAGCAAACCAGGGG | Cas9 | 36.1 | 75.0 | 86.0 | TRUE | TRUE | |
| CTGGGGTGCCTTGGGGATCTGGG | Cas9 | 36.0 | 25.5 | 78.0 | TRUE | TRUE | |
| TGAGAGCAGGGTCGGGGCACGGG | Cas9 | 35.9 | 38.8 | 71.0 | TRUE | TRUE | |
| TTGTTTTCTGCTATTGCCTGTGG | Cas9 | 35.7 | 45.8 | 81.0 | TRUE | TRUE | |
| GGACGGGCACTGGGGTGCCTTGG | Cas9 | 35.6 | 32.6 | 76.0 | TRUE | TRUE | |
| AGGCATCGGAAAATCCACAGAGG | Cas9 | 35.6 | 76.9 | 86.0 | TRUE | TRUE | |
| CCAGTCCGCCCAGGTCCCCTTGG | Cas9 | 35.5 | 42.9 | 74.0 | | TRUE | |
| CCCCAGTGCCCGTCCCTTCCTGG | Cas9 | 35.4 | 31.6 | 83.0 | TRUE | TRUE | |
| CAATGCTCAGAAAACCCCACAGG | Cas9 | 35.4 | 66.2 | 70.0 | TRUE | TRUE | |
| GGAGCTCAGTGCCCTGAAGTTGG | Cas9 | 35.3 | 50.2 | 73.0 | TRUE | TRUE | |
| AAAGAATTTAAGATGCAGGTTGG | Cas9 | 35.1 | 56.0 | 79.0 | TRUE | TRUE | |
| ACTTCAGGGCACTGAGCTCCTGG | Cas9 | 34.7 | 35.4 | 75.0 | TRUE | TRUE | |
| CCTGGGCGGACTGGCCAGGAAGG | Cas9 | 34.7 | 46.5 | 80.0 | TRUE | TRUE | |

**Table S1. Cas9 and Cas12a gRNA sequences and scores.**

| Sequence (including PAM) | Nuclease | Specificity score (MIT) | Efficiency score (Doench '16 for Cas9, DeepCpf1 for Cas12a) | Lindel score | PM | RM | Synthesized as |
|---|---|---|---|---|---|---|---|
| TCTCTTGGAAACCAACTTCAGGG | Cas9 | 34.6 | 45.3 | 78.0 | TRUE | TRUE | |
| GTCCCATTTTCCAAAGGCATCGG | Cas9 | 34.6 | 53.3 | 55.0 | TRUE | TRUE | |
| GCCTTTGGAAAATGGGACTTAGG | Cas9 | 34.6 | 56.5 | 77.0 | TRUE | TRUE | |
| GGTGGCTGCTGTCCTGACAGAGG | Cas9 | 34.6 | 64.4 | 79.0 | TRUE | TRUE | |
| CGGACTGGCCAGGAAGGGACGGG | Cas9 | 34.5 | 49.7 | 81.0 | TRUE | TRUE | |
| ACCTAAGTCCCATTTTCCAAAGG | Cas9 | 34.4 | 60.8 | 81.0 | TRUE | TRUE | |
| TGTTTTCTGCTATTGCCTGTGGG | Cas9 | 34.4 | 44.4 | 83.0 | TRUE | TRUE | |
| CTGATGGAGTAACTGAGCCTGGG | Cas9 | 34.3 | 53.4 | 70.0 | TRUE | TRUE | |
| GGAAAGAGAACTGTCAGAGTGGG | Cas9 | 33.7 | 59.0 | 77.0 | TRUE | TRUE | |
| AGTAACTGAGCCTGGGGGCTTGG | Cas9 | 33.6 | 26.0 | 79.0 | TRUE | TRUE | |
| GATGGAGTAACTGAGCCTGGGGG | Cas9 | 33.6 | 72.6 | 76.0 | TRUE | TRUE | |
| TTTCCGATGCCTTTGGAAAATGG | Cas9 | 33.5 | 30.6 | 81.0 | TRUE | TRUE | |
| GGTCTTGATGAGAGCAGGGTCGG | Cas9 | 33.4 | 52.4 | 81.0 | TRUE | TRUE | |
| ATGTTCCAAGGGGACCTGGGCGG | Cas9 | 33.3 | 67.5 | 82.0 | | TRUE | |
| GATGAACAATCCTTTTCTCTTGG | Cas9 | 33.3 | 34.2 | 85.0 | TRUE | TRUE | |
| CAACAAAGAATTTAAGATGCAGG | Cas9 | 32.9 | 44.5 | 74.0 | TRUE | TRUE | |
| GTGGGTGAATCCAGCCAGGACGG | Cas9 | 32.8 | 64.1 | 80.0 | TRUE | TRUE | |
| TTCAGGGCACTGAGCTCCTGGGG | Cas9 | 32.7 | 51.5 | 70.0 | TRUE | TRUE | |
| CTGCATCTTAAATTCTTTGTTGG | Cas9 | 32.5 | 29.4 | 79.0 | TRUE | TRUE | |
| TTCCGATGCCTTTGGAAAATGGG | Cas9 | 32.5 | 27.7 | 87.0 | TRUE | TRUE | |
| GACATGTTCCAAGGGGACCTGGG | Cas9 | 32.4 | 59.7 | 76.0 | | TRUE | |
| TGGAAAGAGAACTGTCAGAGTGG | Cas9 | 31.8 | 63.0 | 86.0 | TRUE | TRUE | |
| TGATGGAGTAACTGAGCCTGGGG | Cas9 | 31.2 | 59.9 | 61.0 | TRUE | TRUE | |
| CTTCAGGGCACTGAGCTCCTGGG | Cas9 | 31.2 | 44.2 | 75.0 | TRUE | TRUE | |
| TGCTGTCCTGACAGAGGCTTAGG | Cas9 | 31.0 | 34.7 | 73.0 | TRUE | TRUE | |
| TGATAAGAATTTCAAATACTTGG | Cas9 | 29.7 | 41.3 | 76.0 | | TRUE | |
| CTTATTAAATGTTAAAAGACAGG | Cas9 | 29.6 | 45.5 | 81.0 | TRUE | TRUE | |
| GAAGTGGCTCCAGAGAAAAATGG | Cas9 | 29.0 | 43.5 | 77.0 | TRUE | TRUE | |

**Table S1. Cas9 and Cas12a gRNA sequences and scores.**

| Sequence (including PAM) | Nuclease | Specificity score (MIT) | Efficiency score (Doench '16 for Cas9, DeepCpf1 for Cas12a) | Lindel score | PM | RM | Synthesized as |
|---|---|---|---|---|---|---|---|
| ATGGCTGAGGAATGTGTCTCAGG | Cas9 | 28.8 | 52.7 | 78.0 | TRUE | TRUE | |
| TGGGGATCTGGGAGCCTCTGTGG | Cas9 | 28.8 | 50.0 | 82.0 | TRUE | TRUE | |
| GCGGACTGGCCAGGAAGGGACGG | Cas9 | 28.5 | 48.4 | 82.0 | TRUE | TRUE | |
| CAGGCAATAGCAGAAAACAAAGG | Cas9 | 28.5 | 66.9 | 83.0 | TRUE | TRUE | |
| TTGTCTTAACCATTTTTCTCTGG | Cas9 | 28.4 | 34.5 | 78.0 | TRUE | TRUE | |
| AAGTTGGTTTCCAAGAGAAAAGG | Cas9 | 28.3 | 41.7 | 81.0 | TRUE | TRUE | |
| CCCTGCTCTCATCAAGACCAAGG | Cas9 | 28.0 | 66.6 | 64.0 | TRUE | TRUE | |
| CAGAGGCTTAGGGAGGCCCCAGG | Cas9 | 27.2 | 48.9 | 86.0 | TRUE | TRUE | |
| CAGAGGCTCCCAGATCCCCAAGG | Cas9 | 24.8 | 62.4 | 82.0 | TRUE | TRUE | |
| AAGACAGGATATGTTTGAAGTGG | Cas9 | 24.2 | 58.1 | 76.0 | TRUE | TRUE | |
| ACATTTTCTTTAGAATTATGAGG | Cas9 | 22.3 | 53.4 | 71.0 | TRUE | TRUE | |

All possible gRNA sequences at the NHP *IGH* locus identified for Cas9 and Cas12a. Protospacer adjacent motifs (PAM) are included for each sequence. Specificity scores, efficiency scores, and Lindel scores were included where available. Sequences were included for both *Macaca nemestrina* (pigtail macaque, PM), and *M. mulatta* (rhesus macaque, RM). gRNAs were sorted by specificity score first, and the top 5 for each nuclease with homology to both PM and RM were synthesized. NHP_IGH_Cas12a_5 is referred to as the optimal Cas12a gRNA in text.

**Table S2. NHP CD34 purity and yield following enrichment.**

| Animal | Sex | Age (years) | Weight (kg) | Volume of bone marrow draw (mL) | Pre-enrichment count (viable cells) | Pre-enrichment viability | CD34$^+$ fraction count (viable cells) | CD34$^+$ fraction viability | % CD34$^+$ post-enrichment |
|---|---|---|---|---|---|---|---|---|---|
| M03312 | Female | 18.8 | 8.83 | 70.0 | 2.06E+09 | 92.0% | 3.59E+07 | 69.0% | 83.9% |
| Z12214 | Male | 10.1 | 13.34 | 87.5 | 7.28E+08 | 95.0% | 8.72E+06 | 80.5% | 95.2% |
| Z09133 | Female | 13.5 | 8.94 | 90.0 | 1.00E+09 | 91.5% | 8.00E+06 | 85.5% | 97.1% |
| Z14020 | Male | 8.9 | 18.27 | 100.0 | 1.05E+09 | 94.0% | 7.28E+06 | 71.5% | 87.9% |
| Z18090 | Male | 5.7 | 11.86 | 50.0 | 7.75E+08 | 89.0% | 1.82E+07 | 79.0% | 95.8% |
| Z14378 | Female | 8.7 | 8.56 | 75.0 | 5.02E+08 | 95.5% | 7.95E+06 | 78.5% | 79.4% |
| Z16134 | Male | 7.6 | 18.32 | 120.0 | 1.55E+09 | 94.0% | 9.95E+06 | 74.0% | 86.0% |
| Z14122 | Male | 9.8 | 11.86 | 60.0 | 8.86E+08 | 94.0% | 7.08E+06 | 75.0% | 84.3% |
| Z15386 | Female | 8.2 | 7.8 | 70.0 | 2.88E+09 | 90.0% | 3.29E+07 | 87.5% | 91.6% |

Donor information for all *M. nemestrina* animals from which bone marrow aspirates (BMA) were collected for CD34$^+$ enrichment. Viable cell counts were obtained by trypan blue dye exclusion staining pre- and post-enrichment. The percentage of CD34$^+$ cells was measured by flow cytometry, prior to cryopreservation.

**Table S3. Primers for sequencing and template amplification.**

| Primer use | Name | Sequence |
|---|---|---|
| Sanger sequencing | NHP_IGH_F1 | CCACTCTAGAGCCTTTGTTTTCTGC |
| | NHP_IGH_R1 | CTTGGAAACCAACTTCAGGGCACTG |
| | NHP_IGH_F2 | CTGGTCCTCGGGACATGTTCCAAGG |
| | NHP_IGH_R2 | GGCCTCCCTAAGCCTCTGTCAGGAC |
| | NHP_IGH_F3 | ATGCCTTTGGAAAATGGGACTTAGG |
| | NHP_IGH_R3 | GACCCTGCTCTCATCAAGACCAAGG |
| MiSeq library | MiSeq_NHP_IgH_F1 | TCGTCGGCAGCGTCAGATGTGTATAAGAGACAGGACATTCTGCCATTGTGATTACT |
| | MiSeq_NHP_IgH_R1 | GTCTCGTGGGCTCGGAGATGTGTATAAGAGACAGAGGCTCAGTTACTCCATCAG |
| | MiSeq_NHP_IgH_F2 | TCGTCGGCAGCGTCAGATGTGTATAAGAGACAGCTGATGGAGTAACTGAGCCT |
| | MiSeq_NHP_IgH_R2 | GTCTCGTGGGCTCGGAGATGTGTATAAGAGACAGCACCCTTTCTGGACTACCC |
| | MiSeq_NHP_IgH_F3 | TCGTCGGCAGCGTCAGATGTGTATAAGAGACAGGCCTTGGTCTTGATGAGAGC |
| | MiSeq_NHP_IgH_R3 | GTCTCGTGGGCTCGGAGATGTGTATAAGAGACAGAGCACTGTGCTAGTATTTCTTAG |
| PacBio library | M13_IGH_HDR_F10 | /5AmMC6/GTAAAACGACGGCCAGTTGGAAAATGGGACTTAGGTTGG |
| | M13_IGH_HDR_R11 | /5AmMC6/CAGGAAACAGCTATGACGTTCTCTAGCAGGCTTAGGTCT |
| Template amplification | IGH_HDT_F1 | GACATTCTGCCATTGTGATTACT |
| | IGH_HDT_R1 | AGCACTGTGCTAGTATTTCTTAG |
| | IGH_HDT_F2 | GATGGACATTCTGCCATTGTGAT |
| | IGH_HDT_R2 | CTGTGCTAGTATTTCTTAGCTAA |

Primer sequences used for amplifying the NHP *IGH* locus and associated transgene templates by PCR. Sanger sequencing of amplicons was performed in both the forward and reverse primer direction. MiSeq primers include Illumina adapter sequences, PacBio primers include M13 adapters. MiSeq library preparation was performed using primer pair 1 for Cas9 gRNA #1, primer pair 2 for Cas9 gRNA #2-4 and Cas12a gRNA #2, 3 and 5, and primer pair 3 for Cas9 gRNA #5 and Cas12a gRNA #1 and 4. Template amplification was performed using primer pair 1 for CMV-GFP$^{Cas12a}$, and primer pair 2 for VH4-GFP$^{Cas12a}$ and VH4-10-1074$^{Cas12a}$. 5AmMC6: 5’ Amino Modifier C6 (IDT).

**Table S4. Primatized MISTRG and MISTRG6 cohort conditions.**

| Cohort | Strain | NHP Donor(s) | Condition | Template | n (injected) | n (deceased) | $CD34^+$ cell dose |
|---|---|---|---|---|---|---|---|
| 1 | MISTRG | Z14020 | CMV-GFP | dsDNA | 6 | 1 | 2.54E+04 |
| 2 | MISTRG | Z09133; Z14020 | VH4-GFP | dsDNA | 8 | 0 | 1.39E+05 |
| | | | Control | NA | 7 | 1 | 1.71E+05 |
| 3 | MISTRG | Z16134; Z18090 | VH4-10-1074 | dsDNA | 5 | 0 | 5.00E+05 |
| 4 | MISTRG | Z15386; Z16134 | VH4-10-1074 | AAV6.2 | 2 | 0 | 5.00E+05 |
| | | | Control | NA | 2 | 1 | 5.00E+05 |
| 5 | MISTRG6 | Z15386; Z16134 | VH4-10-1074 | dsDNA | 4 | 1 | 5.00E+05 |
| | | | VH4-10-1074 | AAV6.2 | 4 | 1 | 5.00E+05 |
| | | | Control | | 2 | 1 | 5.00E+05 |

MISTRG or MISTRG6 mice born on the same day were grouped into cohorts, each of which was injected with NHP HSPC from different donors and conditions as shown here. Results from cohort #1 were not included in the main text due to low engraftment and lack of GFP expression. All deceased MISTRG mice died prior to week 8 from causes seemingly unrelated to injection (no apparent signs of GVHD). Deceased MISTRG6 mice showed clear signs of GVHD. Mice that died prior to week 8 were not included in our analysis.

**Table S5. Antibodies for flow cytometry on MISTRG samples.**

| Marker | Fluor | Clone | Dilution | Peripheral Blood Panel | Necropsy Panel |
|---|---|---|---|---|---|
| **mCD45** | V500 | 30-F11 | 1/100 | TRUE | TRUE |
| **pCD45** | V450 | D058-1283 | 1/50 | TRUE | TRUE |
| **Anti-Strep Tag II** | FITC | 5A9F9 | 1/50 | TRUE | TRUE |
| **pCD138** | BUV737 | MI15 | 1/50 | TRUE | TRUE |
| **pCD20** | APC | 2H7 | 1/50 | TRUE | |
| **pCD3** | BV650 | SP34-2 | 1/50 | TRUE | |
| **pCD14** | PE-Cy7 | 61D3 | 1/50 | TRUE | |
| **pCD34** | PE | 563 | 1/50 | | TRUE |
| **pCD19** | PE-eFluor610 | HIB-19 | 1/50 | | TRUE |
| **pCD4** | PE-Cy7 | SK3 | 1/50 | | TRUE |
| **pCD8a** | AF 647 | RPA-T8 | 1/50 | | TRUE |

Anti-mouse (m) and anti-primate (p) antibodies used for flow cytometry analysis in MISTRG studies. Cross-reactivity of clones in target species were confirmed on the Non-Human Primate Reagent Resource (NHPRR).

**Table S6. Antibodies for flow cytometry on MISTRG6 samples.**

| Marker | Fluor | Clone | Dilution |
|---|---|---|---|
| **mCD45** | V500 | 30-F11 | 1/100 |
| **pCD45** | V450 | D058-1283 | 1/50 |
| **Anti-Strep Tag II** | FITC | 5A9F9 | 1/50 |
| **pCD138** | BUV737 | MI15 | 1/50 |
| **pCD31** | BV650 | WM59 | 1/50 |
| **pIgG** | BV786 | G18-145 | 1/50 |
| **pCD20** | BUV395 | 2H7 | 1/50 |
| **pCD34** | PE | 563 | 1/50 |
| **pCD90** | PerCP-Cy5.5 | 5E10 | 1/50 |
| **pCD4** | PE-Cy7 | SK3 | 1/50 |
| **pCD8a** | AF 647 | RPA-T8 | 1/50 |

Anti-mouse (m) and anti-primate (p) antibodies used for flow cytometry analysis in MISTRG6 studies. Cross-reactivity of clones in target species were confirmed on the Non-Human Primate Reagent Resource (NHPRR). Peripheral blood and necropsy panels were identical.

### SUPPLEMENTARY FIGURES

**Figure S1.**

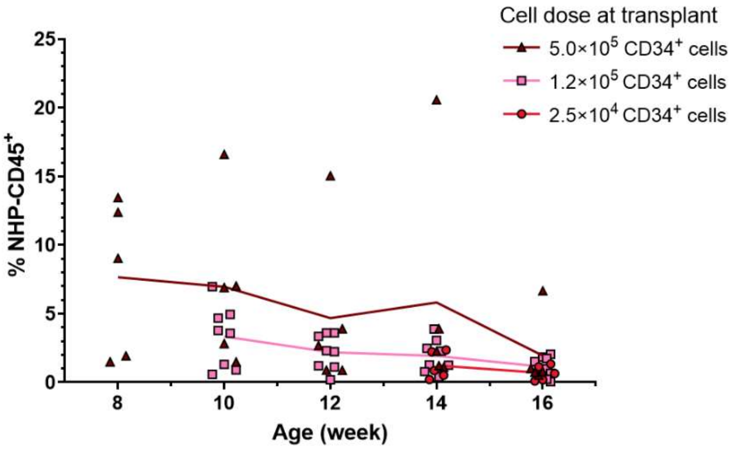


**Figure S1. NHP HSPC engraftment in MISTRG mice at escalating cell doses.** Engraftment levels in peripheral blood over time as measured by flow cytometry on NHP-CD45$^+$ cells in mice injected with different doses of electroporated HSPC. n = 5 mice, n = 1 NHP donor ($2.5 \times 10^4$ cell dose group). n = 8 mice, n = 2 NHP donors ($1.2 \times 10^5$ cell dose group). n = 5 mice, n = 2 NHP donors ($5.0 \times 10^5$ cell dose group).

**Figure S2.**

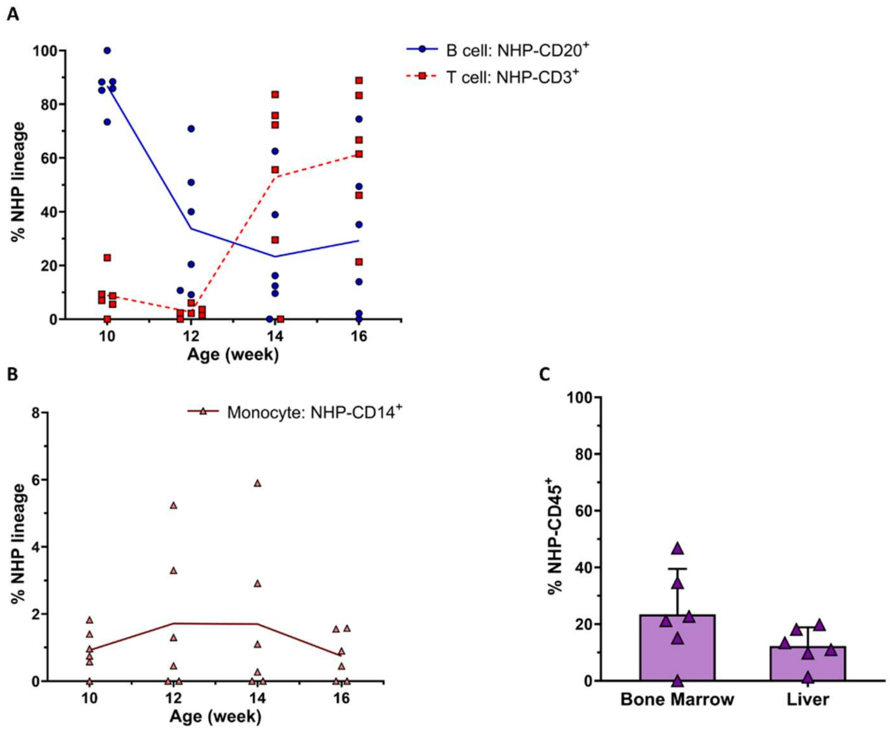


**Figure S2. Control NHP HSPC engraft and differentiate into mature cell types in MISTRG model.** (A, B) Multilineage engraftment in peripheral blood over time as measured by frequency of B cells (CD3$^-$ CD20$^+$), T cells (CD3$^+$ CD20$^-$), and monocytes (CD3$^-$ CD20$^-$ CD14$^+$) in NHP-derived populations. (C) Engraftment levels in the bone marrow (femur) and the liver at necropsy, performed at week 20. n = 6 mice, n = 2 NHP donors (control group, week 18 omitted for low engraftment levels, A-B). Error bars represent SD (C).

**Figure S3.**

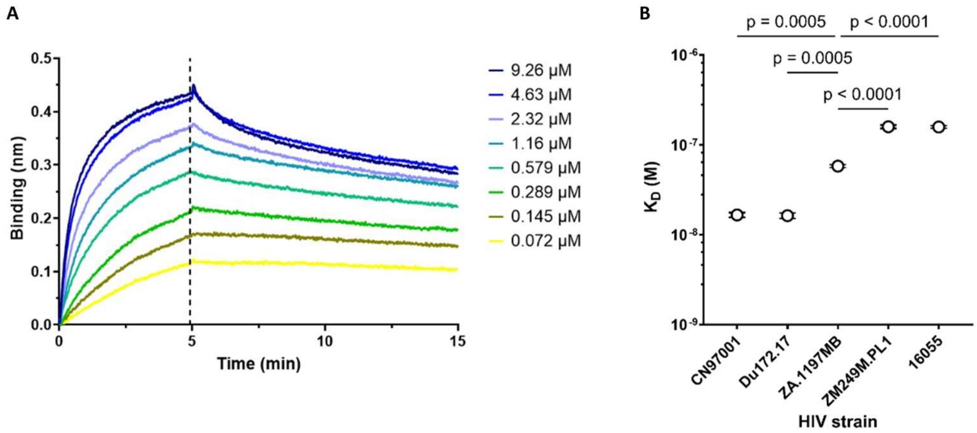


**Figure S3. Recombinant 10-1074 antibody binds to HIV antigen.** (A) Binding of 1 µM gp120 monomer from HIV CN97001 to recombinant 10-1074 at varying antibody concentrations as measured by wavelength shift using Bio-Layer Interferometry (BLI). Binding and dissociation phase, separated by dotted line, are shown here. Samples were aligned at beginning of binding phase. Lines represent mean. (B) Dissociation constant ($K_D$) calculated from BLI measurements for binding of recombinant 10-1074 to gp120 monomer from different HIV strains. Data are presented as mean ± SD. Significance was calculated with an ordinary one-way ANOVA. n = 2 technical replicates.

**Figure S4.**

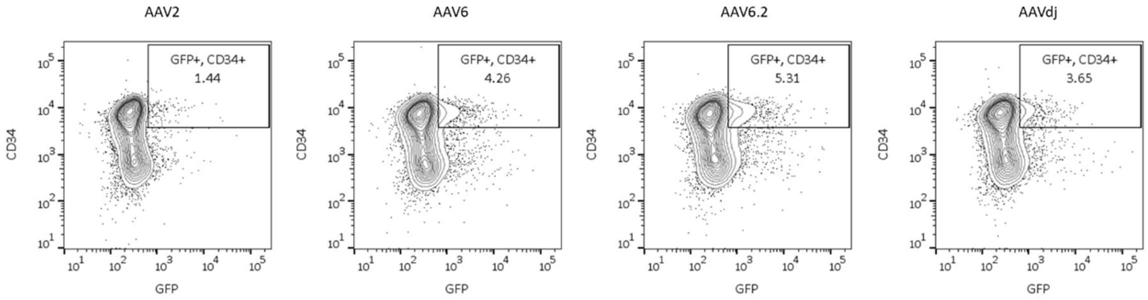


**Figure S4. AAV6.2 transduces NHP HSPC and leads to GFP expression.** $CD34^+$ cells were transduced with AAV from 4 different serotypes containing a CMV-mediated GFP reporter at an MOI of $6.67 \times 10^5$. GFP expression was measured by flow cytometry 24 hours later. Transduction at an MOI of $2.40 \times 10^4$ did not produce significant GFP expression.

**Figure S5.**

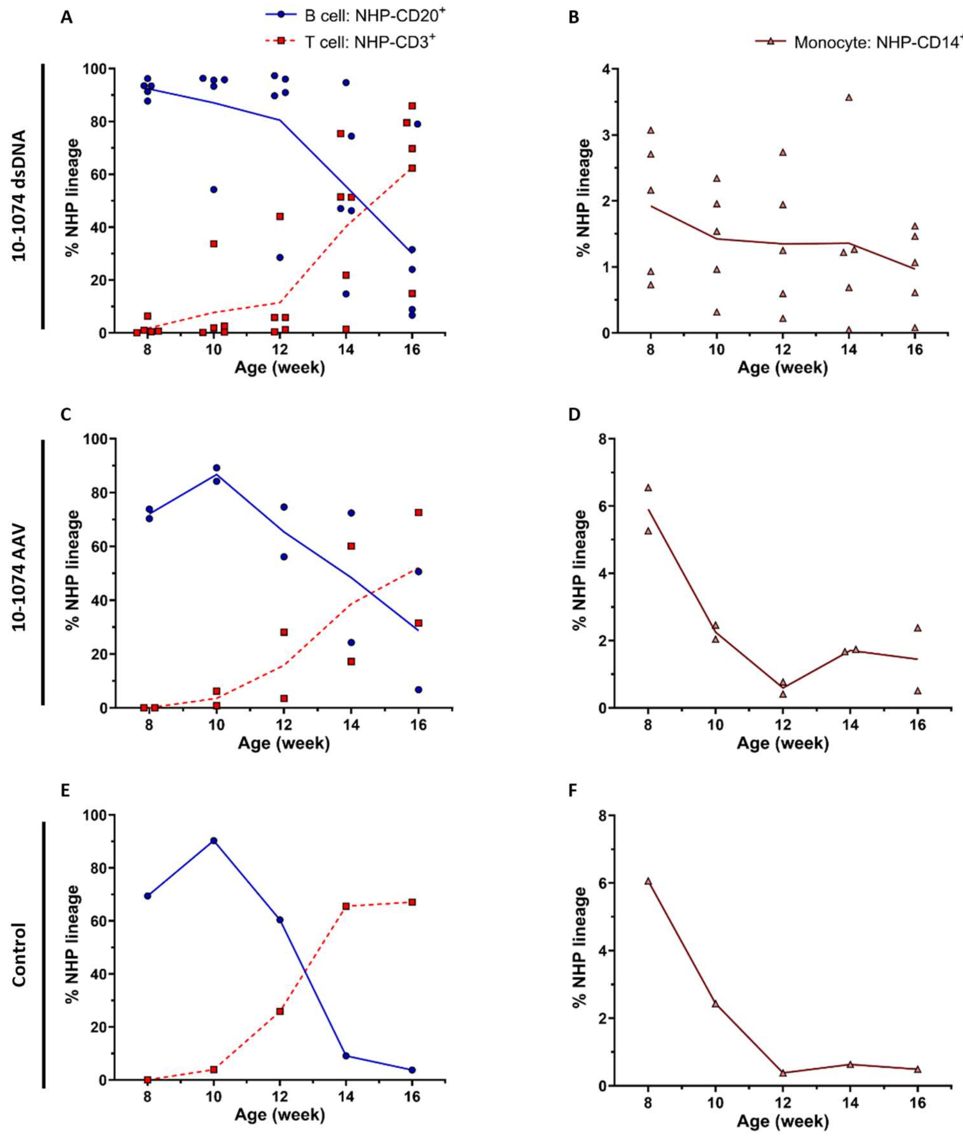


**Figure S5. Hematopoietic lineages form in MISTRG mice injected with HSPC with or without different DNA templates.** Multilineage engraftment in peripheral blood over time as measured by frequency of B cells ($CD3^-$ $CD20^+$), T cells ($CD3^+$ $CD20^-$), and monocytes ($CD3^-$ $CD20^-$ $CD14^+$) in NHP-derived populations for mice receiving NHP HSPC electroporated with dsDNA (A, B) or AAV (C, D), as compared to mice receiving control NHP HSPC (E, F).

**Figure S6.**

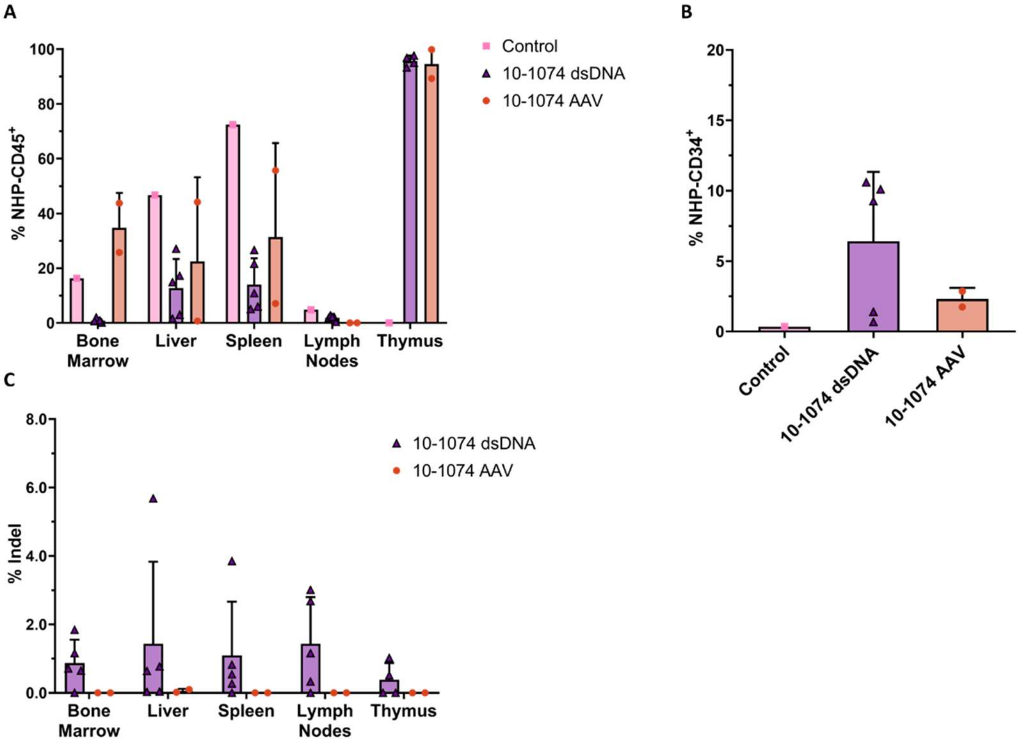


**Figure S6. Engraftment and editing levels in various tissues at necropsy for MISTRG mice receiving different DNA templates.** (A) Fraction of NHP-CD45⁺ in single cell suspensions from the bone marrow (femur), liver, spleen, lymph nodes, and thymus at necropsy, performed week 18. (B) Fraction of NHP lineage that is CD34⁺ in the bone marrow. (C) Editing levels at the NHP *IGH* locus in the bone marrow, liver, spleen, lymph nodes, and thymus, as measured by fraction of MiSeq reads containing indels. n = 5 mice, n = 2 NHP donors (10-1074 dsDNA group). n = 2 mice, n = 2 NHP donors (10-1074 AAV group). n = 1 mouse, n = 2 NHP donors (control group). Error bars represent SD.

**Figure S7.**

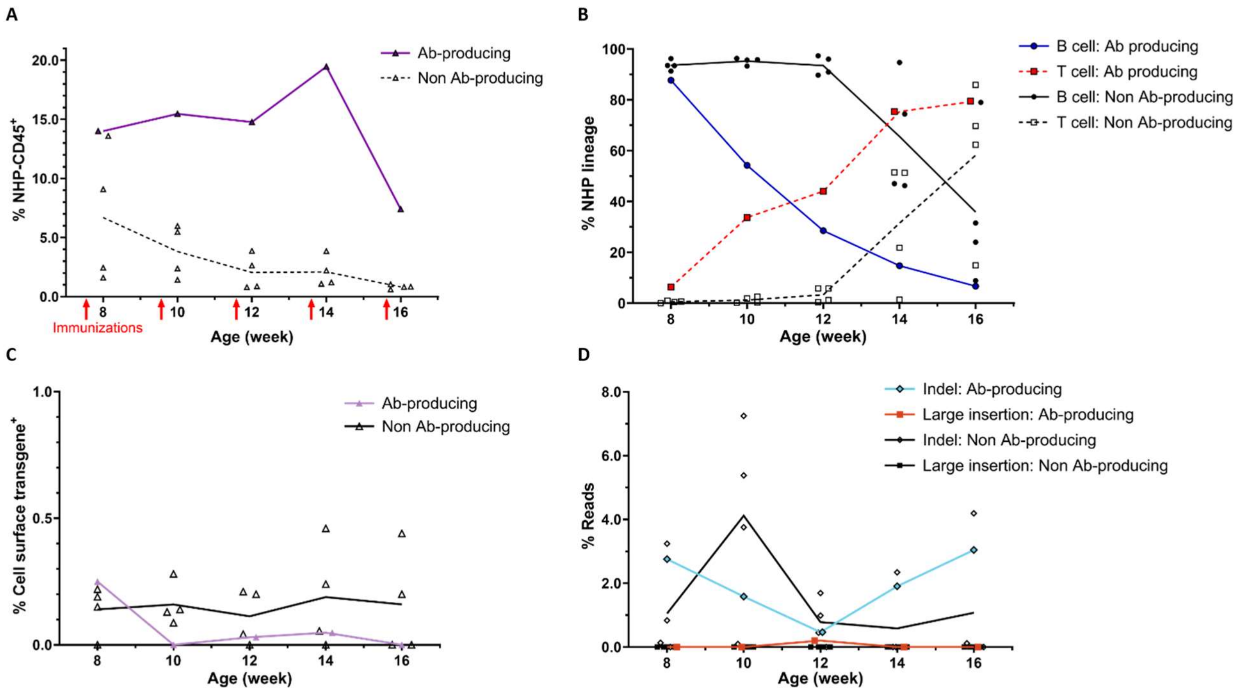


**Figure S7. Engraftment, cell lineage, and editing in antibody-producing MISTRG mouse.** (A) Engraftment levels in peripheral blood over time as measured by flow cytometry on NHP-CD45$^+$ cells in 10-1074-producing mouse and non 10-1074-producing mice. (B) Lymphocyte populations in peripheral blood over time as measured by frequency of B cells (CD3$^-$ CD20$^+$) and T cells (CD3$^+$ CD20$^-$) in NHP-derived populations for 10-1074-producing mouse and non 10-1074-producing mice. (C) Cell surface expression of transgene in peripheral blood as measured by flow cytometry with antibody against its linker for 10-1074-producing mouse and non 10-1074-producing mice. (D) Editing levels at the NHP *IGH* locus in peripheral blood of 10-1074-producing mouse and non 10-1074-producing mice. Large insertions representing insertions >8 bp. The MISTRG mouse (n = 1 of 5, 20%) with detectable anti-HIV antibody titers has been singled out in each graph.

**Figure S8.**

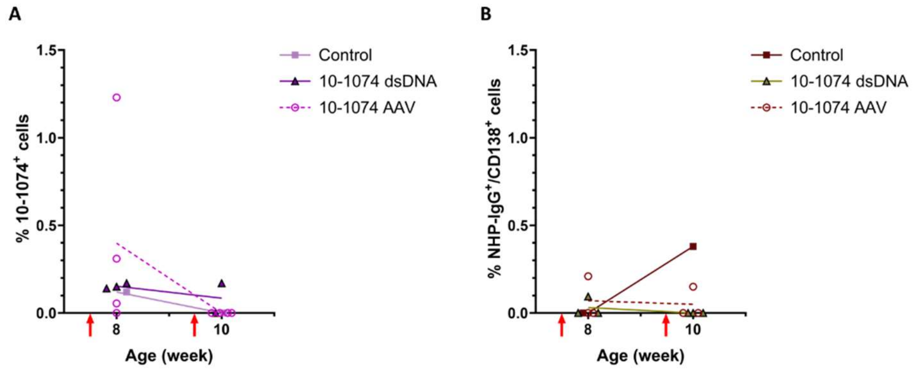


**Figure S8. 10-1074-expressing and IgG-producing cells in circulation in MISTRG6.** (A) Cell surface expression of 10-1074 transgene in peripheral blood as measured by flow cytometry with antibody against its linker. (B) IgG-producing cells as measured by frequency of NHP-IgG$^+$/CD138$^+$ cells in NHP-derived populations (NHP-CD45$^+$). Red lines in represent immunizations.

**Figure S9.**

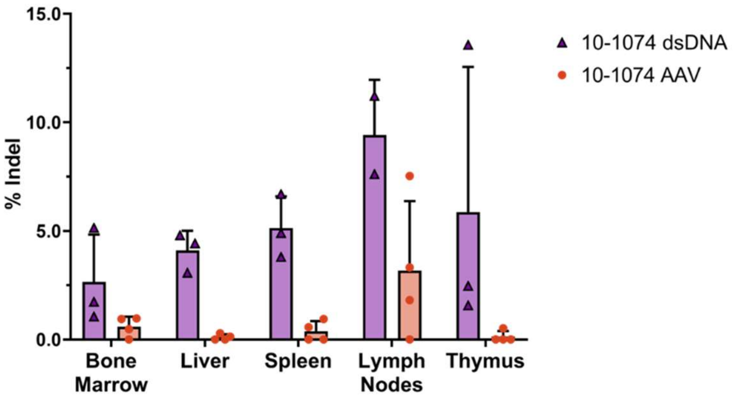


**Figure S9. Editing levels in various tissues at necropsy for MISTRG6 mice.** Editing levels at the NHP *IGH* locus in the bone marrow, liver, spleen, lymph nodes, and thymus, as measured by fraction of MiSeq reads containing indels.

**Figure S10.**

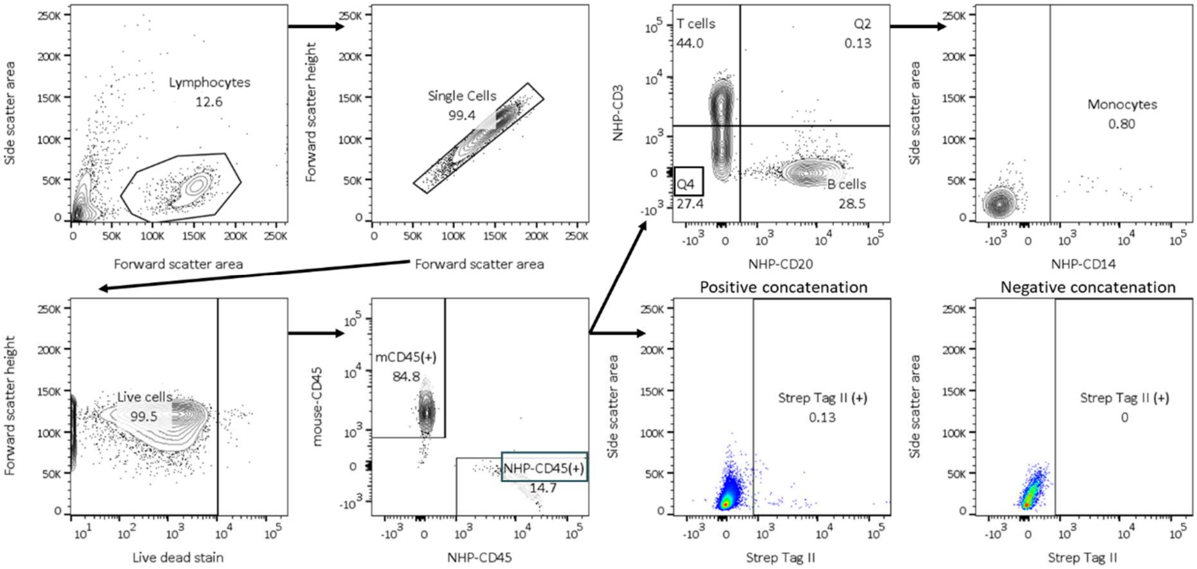


**Figure S10. Flow cytometry gating strategy for MISTRG samples.** Example of gating strategy for flow cytometry of peripheral blood from antibody-producing mouse at week 12. Compensation was calculated for every fluorophore and each sample. For CD34 enriched samples, live cells were only gated on CD34 levels. For necropsy samples, B cells were gated as NHP-CD19$^+$/CD4$^-$ /CD8a$^-$. Strep Tag II gates shown as concatenation of positive or negative samples due to low event counts.

**Figure S11.**

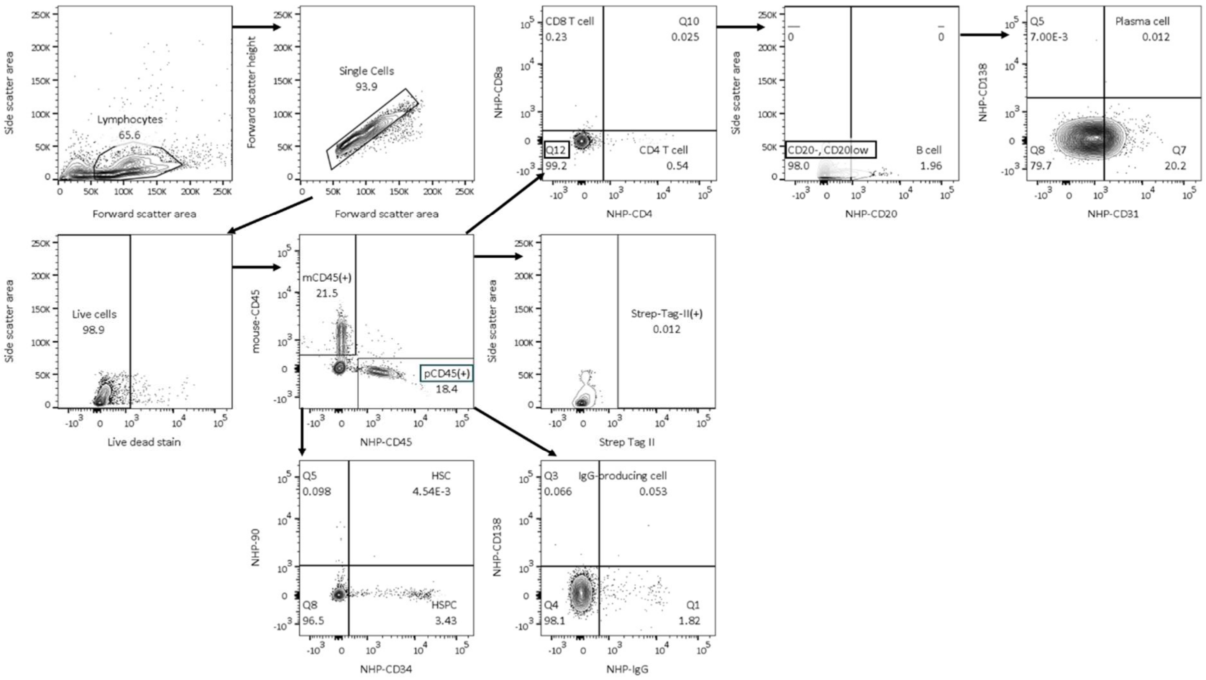


**Figure S11. Flow cytometry gating strategy for MISTRG6 samples.** Example of gating strategy for flow cytometry of bone marrow from mouse injected with HSPC electroporated with dsDNA. Peripheral blood samples were gated using the same strategy. Compensation was calculated for every fluorophore and each sample.

**Sequence S1. Template knock-in at *M. nemestrina IGH***

**Legend:** Upstream homology arm (PAM blocking mutation) : Site for barcode integration : VH4 promoter including Kozak sequence : Recombinant 10-1074 including splice site : Site for barcode integration : Downstream homology arm (Mutation for synthesis) : Reference sequence

TTGGAAAATGGGACTTAGGTTGGATGTGTCTGATGGAGTAACTGAGCCTGGGGGCTTGGG
GAGCCACATTTGGACGAGATGCCAGAGCAAACCAGGGGTCTTAGTGATGGCTGAGGAATG
TGTCTCAGGTGCGGTGTCTGTAGGACTGCAAGATCGCTGCACACACAGCGAATCATGAAA
CATTATCTTTAGAATTATGAGGGATCCTCGGACGAATTCCACACTCTCCGGGACAGATATAT
TCCCTCTAACCATGATGGATATTCTGAATTACAATAAACATTGCACGGATGTAGGTTTATAT
AAATTCACTGTGACGAATATATTTAGCTGCTGCCCTAATGTTTTGAATGGAGATTTGACAAT
TTAGATAACCTGGATGTTTGGTTGGTTTCATATAAATCTTCAAGGGTACAACAGCATTGAAC
CTATTCCAAAATCTATCCCTGATCCATGATCATACTCATCTCCCAGACCAGCTCCTTCAGCA
CATCTCCCTACCTGGAAGAAGAGGAGTCTGGGCTTGGTGAGGGGAGGCCCCAGGAAGAG
AACTGGGTTCTCAGAGGGCACAGCCAGCATCCTCCTCTCAGGGTGAGCCCCAAAGACTGA
GGCCTCCCTCATCCCTTTTCACCTCTCCATACAAAGGCACCACCCACATGCAAATCCTCAC
TTAGGCACCCACAGGAAACCACCACACATTTCCTTAAATTCAGGGTCCAGCTCACATGGGA
AATGCTCTCTGAGAGTCACGGACCTCCTGTGCAAGAACATGAAGCACCTGTGGTTCTTCCT
CCTCCTGGTGGCAGCTCCCAGATGTGAGTGTAGCTACGTTCGACCTCTCAGCGTTGCATT
GGGAGAAACTGCTCGGATATCCTGCGGCAGGCAGGCCTTGGGGAGTCGGGCAGTACAAT
GGTACCAACATAGACCAGGCCAGGCTCCAATCTTGTTGATTTATAATAATCAGGATAGGCC
AAGCGGAATCCCCGAGAGGTTTTCTGGCACACCAGATATAAACTTTGGGACCCGAGCAAC
ACTCACTATCTCTGGGGTCGAGGCCGGAGACGAGGCTGACTACTATTGCCATATGTGGGA
TTCAAGGTCCGGTTTCTCTTGGTCCTTTGGAGGTGCCACACGACTCACAGTGCTGGGCGG
GCAGCCAAAAGCATCTCCTTCCGTAACACTCTTTCCTCCATCATCTGAAGAATTGCAAGCC
AATAAGGCTACCCTCGTTTGCCTCATATCTGATTTTTATCCAGGCGCAGTCGAAGTGGCCT

GGAAAGCTGATGGGTCTGCAGTCAATGCAGGTGTCGAGACAACAAAGCCTTCAAAGCAGTCCAACAACAAGTACGCCGCCTCTTCCTACTTGTCTCTCACATCTGATCAATGGAAAAGCCATAAGTCTTACAGCTGCCAGGTGACTCACGAAGGGAGCACTGTAGAGAAGACCGTCGCTCCCGCTGAATGTTCTGGAGGCAGTTCTGGATCTGGGTCCGGAAGCAACTGGTCTCATCCACAGTTCGAAAAAGGAGGGGGTGGATCAAATTGGTCACACCCACAGTTTGAAAAAGGAGGTGGAGGTTCCAATTGGAGCCACCCACAATTCGAGAAAGGCTCAGGAGGCGGAGGGAGCGCAGGTGGGCAGGTGCAGTTGCAAGAGTCCGGTCCTGGACTCGTTAAGCCATCTGAAACTCTTAGCGTCACCTGCTCTGTCTCCGGAGACTCCATGAACAACTACTACTGGACTTGGATTCGGCAATCACCCGGCAAGGGACTTGAATGGATCGGCTATATCTCAGACCGCGAATCTGCAACATACAATCCATCTCTCAATAGCAGAGTTGTTATTTCACGAGATACTTCAAAAAATCAGTTGTCCTTGAAGCTTAATTCAGTAACCCCCGCTGACACAGCCGTGTACTATTGTGCTACTGCCAGACGCGGACAACGGATCTATGGTGTTGTAAGTTTCGGGGAGTTTTTTACTATTATTCAATGGATGTCTGGGGGAAGGGAACTACTGTCACAGTGAGCAGCGGTGAGTCCTCACAACCTCTCTCCTGCTTTAACTCTGAGGGGTTTTGCTGCATTTTTGGGGGAAAACAAGTGTGGCGGCCGCTACGTAACTAGTGTGCGCTGTGTGCCAACCTGCATCTTAAATTCTTTGTTGGCTGGAAAGAGAACTGTCAGAGTGGGTGAATCCAGCCAGGACGGATGCGTAGCCTTGGTCTTGATGAGAGCAGGGTCGGGGCACGGGTAGTCCAGAAAGGGTGGCTGCTGTCCTGACAGAGGCTTAGGGAGGCCACAGGAGCTCAGTGCCCTGAAGTTGGTTTCCAAGAGAAAAGGATTGTTCATCTTAAGAGGCGTGCTTATTAAATGTTAAAAGACAGGATATGTTTGAAGTGGCTCCAGAGAAAAATGGTTAAGACAATTATGACTTAAAAATGTGAAAGATTTTGAAGTATATTAATTTTTTTAACTGTCTAAGTATTTGAAATTCTTATCATTTGATTAACACCCATGAGTGGTATGTCTCTGGAATTAGGGCCAAAGTAAGTTTAGCTAAGAAATACTAGCACAGTGCTGTCGGCTCTGATGCAGGACTGAGTTTTGAGCATCATAAATCAAGTTTATTTTTTTAATTAATTGAGTGAAGCTGGGAGCGGATGATGAGTTAGAGTCAAGATGGCTGCATGGGGGTCTTGGGCACCCACAGCAGGTGGCAGGAAGCAGGTCACCGCAAGAGTCTATTTTAGGAAGCAAAAAAACACAATTGGTAAATTTATCACTTCTGGTTGTGAAGAGGTGGTTTCGCCTGGGCCCAGATCTGAAAGTGCTCTACTGAGCAAAACAACA

CCTGGACGATTTGCGTTTCTAAAATAAGGCAAGGCTGACCGAAACTGGAAAGGCTCTTTTT
TTTTTAACTATCAGAATTTCATTTCCAATCTTAGCTTATCAACTGCTAGTTTGTGCAAACAGG
ATATCAACTTCTAAACTGCATTCATTTTTACAGTAAGATGTTTAAGAAATTAAAGAGTCTTAG
GGAGAGTTTATGACTGCATTCAAAAAACTTTTAAAATTAGTCTGTTATCCCTTCATGTGATAA
TTAATCTCAAACACGTTTTCAATACCTCAGAGCATTATTTTTATAATAACTATGTTCACAATCT
TTTTAGGTTAACTCGTTTTCTCTTTGTGATTAAGGAGAAACACTTTGATATTCTGATAGAGTG
GCCTTCATTTTAGTTGTTTTTCAAGACCACTTTTCAACTACTCACTTTAGGATAAGTTTTAGG
TAAAATATGCATCATTATCCTGAATTATTTCAGTTAAACATGTTAGTTTGTGGCTTAAGAGAA
AACTCAGTCAGATAGTGCTGAAGACAGGACTGTGGAGACACCTTAGAAGGACGGGTTCTG
TTCGGAATCACCGATTCGGCTTCAGCCAGACTGGCCTAGCGGAGGCTCTGGGAGGCTGC
CTGCCAGGCCCGGGCTGGGCTTTGGGTCTCCCCGGACTACCCAGAGCTGGGATGCGTGG
CTTCTGCCACCAGACCAACTGGCTGCTCAGGACCCAGCCCTTGTTAATGGACTTGGAGGA
ATGATTCCATGCCAAAGCCTTGCAAGGCTCGCAGTGACTACACACACCGACATGGTAAGA
GACAGGCAGCCGCCGCTGCTGCATTCGCTTCTCTTAAACCTTTGTATTTGATGTCTTATTTG
CACTAGAAGGGGAATTGGTCTTTGCTTGATGAAGAGCAGGAGACTCATTCGTGTGAGTCTT
TTGACTGACCATTGTCTGGGTCACTCCCATTTAACTTTCCCTAAAGCCCATTTGAAGGAGAG
GTCACCCGAGCTCTTTCGCAACCTCTGAATGCGGATGGCATGAGTCATGATGCTTCAGCC
CCACTGGCATCGCCCTTGTCTAAATCATTGACTTAGGTCATCGCGCCCTTGAAAGTAGCCC
ATGCCTTCCAAAGCGATTTGCGGTAAATGGCAGAATTTTAAGTGGCAAATTCAGATAAAATG
CATTTCTTGGTTGTTTCCAATGATGACTGTTATCTAGAGGGAATTTAAAGGCAGGGGTTTAT
TTCAGACTCAGAAGGGAGGAGATGCTCTGAGAAGGTGGAGGCTTTGAGCATTTCAGTACC
CTCCTCTCGGTGCAGAAGCTATGCTGCCGATTCTAGAGCAAGGGGAGCTCCTCATTTTTAT
CACAGTACAGGCTCCTAAATTCTCGGTCTAATTCTGAAGATGTTCTAATAACTTTAAAGCAG
CAAAGAAATATTCCACCCAGGTAGTGGAGGGTGGTAATGATTGGTACTGCTTTGGAACCAA
AACCCAGGTGGTGCTAGGGCAGGACTGCAGGAAACTGGGGTATCAAGTAGAGGGAGACA
AAAGATGAAAGCCGGCCTGTGCAGGAACCTAGCAATGAGACGGCCTTAGCTGAGACAAGC

AGGTCTGGTGGGCTGACCGTTTCCAGCCATGACAACTCCATCCAGCTTTCAGAAATGGCCT
TGGATGGGCAAAACTGACCTAAGCTGACCTAGACTAAACAAGGCTGAACTGCGCTGAGCT
GAGCTGGGTTGAGCTGAGC
